# APOE4 affects basal and NMDAR mediated protein synthesis in neurons by perturbing calcium homeostasis

**DOI:** 10.1101/2020.12.10.418772

**Authors:** Sarayu Ramakrishna, Vishwaja Jhaveri, Sabine C Konings, Sumita Chakraborty, Bjørn Holst, Benjamin Schmid, Gunnar K Gouras, Kristine K Freude, Ravi S Muddashetty

**Affiliations:** Institute for Stem Cell Science and Regenerative Medicine, Bangalore, India; The University of Trans-Disciplinary Health Sciences and Technology (TDU), Bangalore, India; Experimental Dementia Research Unit, Department of Experimental Medical Science, Lund University, 221 84 Lund, Sweden; Bioneer A/S, Kogle Alle 2, 2970 Hørsholm, Denmark; Department of Veterinary and Animal Sciences, Group of Stem Cell Models for Studies of Neurodegenerative Diseases, Section for Pathobiological Sciences, Faculty of Health and Medical Sciences, University of Copenhagen, Denmark

## Abstract

Apolipoprotein E (APOE), one of the primary lipoproteins in the brain has three isoforms in humans – APOE2, APOE3, and APOE4. APOE4 is the most well-established risk factor increasing the pre-disposition for Alzheimer’s disease. The presence of the APOE4 allele alone is shown to cause synaptic defects in neurons and recent studies have identified multiple pathways directly influenced by APOE4. However, the mechanisms underlying APOE4 induced synaptic dysfunction remain elusive. Here, we report that the acute exposure of primary cortical neurons to APOE4 leads to a significant decrease in global protein synthesis. APOE4 treatment also abrogates the NMDA mediated translation response indicating an impairment of synaptic signaling. Importantly, we demonstrate that both APOE3 and APOE4 generate a distinct translation response which is closely linked to their respective calcium signature. Acute exposure to APOE3 causes a short burst of calcium through NMDARs in neurons leading to an initial decrease in protein synthesis which quickly recovers. Contrarily, APOE4 leads to a sustained increase in calcium levels by activating both NMDARs and L-VGCCs, thereby causing sustained translation inhibition through eEF2 phosphorylation, which in turn disrupts NMDAR response. Thus, we show that APOE4 affects basal and activity mediated protein synthesis response in neurons by affecting calcium homeostasis. We propose this as a possible mechanism to explain the synaptic dysfunction caused by APOE4.

**Highlights / Summary:**

- APOE3 treatment causes a short burst of calcium through NMDARs, leading to an acute increase in eEF2 phosphorylation which eventually recovers to basal levels.
- Global translation follows a similar temporal profile of initial inhibition followed by recovery in APOE3 treated neurons, thus unaffecting the NMDA mediated translation response.
- APOE4 treatment activates both NMDARs and L-VGCCs leading to a marked elevation in calcium levels, thus causing sustained increase in eEF2 phosphorylation as well as global translation inhibition.
- Hence, the NMDA mediated response is perturbed, potentially causing a stress-related phenotype in APOE4 treated neurons.
- Thus, different calcium signatures and sources lead to distinct temporal profiles of translation.

## Introduction

Alzheimer’s disease (AD), the most common cause of dementia, is an irreversible progressive neurodegenerative disorder that leads to loss of memory and cognition^1^. Pathologically, AD is characterized by brain atrophy along with an accumulation of Aβ plaques and neurofibrillary tangles^1^. But the most robust correlation for the severity of dementia and staging of AD is with the extent of synapse loss^2^. It is now well established that synapse loss is an early feature and a structural correlate of AD which occurs pre-symptomatically^2^. Synapses are the sites where neuronal activity is interpreted, making them the fundamental functional units of learning and memory. Various stimuli can cause changes in the synaptic function, ultimately bringing about structural changes like induction, pruning, enlargement of spines^3,4^. The re-modeling and maintenance of these structural and functional changes of synapses require the synthesis of new proteins which are tightly regulated spatio-temporally^3,4^. Thus, activity mediated synthesis of new proteins and changes in the proteome play an important role in driving synaptic plasticity^3–5^.

The importance of translation regulation and its link to cognitive deficits is well established in the context of neurodevelopment^5,6^, whereas this link is relatively less studied in neurodegeneration. Multiple studies have shown the dysregulation of protein synthesis and its machinery in AD. The changes in levels, activity, and post-translational modifications of translation components like eIF2α, eEF1A, eIF4E, p70 RPS6 kinase 1 have been previously reported in AD ^7–11^. The altered expression of ribosomal RNA and mRNAs coding for ribosomal proteins are shown to occur before the development of AD symptoms^12–14^. Recent studies using non-canonical amino acid tagging have shown the dysregulation of de-novo protein synthesis in AD^15,16^ and other forms of taupathy as well^17,18^. Along with basal translation, defects in synaptic activity mediated signaling and translation is also reported to occur pre-symptomatically in AD ^7,8,16,19^. These studies establish the defect in protein synthesis machinery and translation regulation in AD. However, the consequence of the translation defects on dysregulation of synaptic signaling is not explored and is likely to be an important early event in AD pathology.

More than 90% of AD cases are sporadic with no known familial mutations in APP or PSEN genes^20^. Beyond the familial mutations, many genetic factors significantly increase the risk of AD^20^. Apolipoprotein isoform ε4 (APOE4) is the most well-established genetic risk factor for AD (both familial and sporadic) which increases the frequency of AD occurrence and decreases the age of onset significantly^21,22^. Unlike its other isoforms ε2 (APOE2) and ε3 (APOE3), APOE4 is shown to increase the predisposition to AD by affecting the clearance of Aβ^21,22^. However, many studies have reported that the presence of the APOE4 allele can cause synaptic defects independently. Reduction of neurite outgrowth, dendritic complexity, spine density, and loss of synaptic proteins is well reported in APOE4 mice models^23–27^. Consequently, APOE4 mice of both younger and older age groups are reported to show defects in spatial learning and memory^23^. This supports the idea that APOE4 could have an impact on cognitive processes from early in life. In support of this, many studies have drawn correlations between APOE genotype and cognition in humans as well ^28–31^. APOE4 is also reported to interfere with glutamate receptor signaling pathways^32^, especially downstream of NMDARs^33–35^. This is further supported by the studies which have shown the cross-talk between APOE receptors and NMDARs^33–35^, indicating that APOE receptors are present as a part of the post-synaptic density complex^36^. APOE, upon binding to its cognate receptor, not only performs the classical function of lipid transport, but it is also known to activate multiple signaling pathways. Several studies have highlighted the key signaling differences between the APOE isoforms as well^37–39^.

Though APOE4 is indicated to be one of the key factors influencing synaptic loss and cognitive defects in AD, the molecular mechanisms behind this are still unclear. Considering the importance of protein synthesis in synaptic functioning and its relevance to the synaptic loss in AD, translation regulation becomes an important aspect in the context of APOE4 mediated defects as well. Hence, we studied the effect of APOE4 on basal and synaptic activity mediated translation in neurons. We show that APOE4 inhibits basal and NMDAR mediated protein synthesis in neurons by increasing eEF2 phosphorylation. This translation response is linked to the increased calcium influx caused by APOE4 by activating NMDARs and L-VGCCs. Thus, the dysregulation of calcium homeostasis by APOE4 leads to its impaired protein synthesis response.

## Results

### APOE4 causes a reduction of protein synthesis in neurons

Activity mediated protein synthesis is a dynamic process which impacts both short-term and long-term aspects of synaptic function and plasticity^3^. APOE4 was previously shown to disrupt many synaptic functions^27,32,40,41^, but its impact on protein synthesis in neurons has not been studied extensively. To study the effect of APOE4 on global and activity mediated protein synthesis in neurons, we used cultured primary cortical neurons (DIV 15) from Sprague-Dawley rats as our model system. We used conditioned or secreted media from human induced pluripotent cells (hiPSCs) as one of the sources of APOE. iPSC lines of different APOE genotypes (APOE3/3, APOE4/4, and APOE KO) were generated by CRISPR-Cas9 based editing of the original APOE3/4 iPSC line^42^. These iPSCs were characterized for the expression of the pluripotency markers (OCT4 and Nanog) and normal karyotype profile (**Fig S1A and S1B**). When the iPSCs reached 50% confluency, they were switched from stem cell medium to neurobasal medium. After 48 hours, the conditioned neurobasal media containing APOE protein was collected from the stem cells. The iPSCs express a significant amount of APOE protein (**Fig S1C)** and the APOE secreted by the iPSCs into the neurobasal media was stable for at least 48 hours (**Fig S1D**). The average concentration of APOE secreted in the conditioned neurobasal media was 0.2-0.3μg/ml as measured by ELISA (**Fig S1E**).

The readouts used for global protein synthesis were - phosphorylation status of eukaryotic translation elongation factor eEF2 (increase in eEF2 phosphorylation implies translation inhibition) and Fluorescent Non-Canonical Amino-acid Tagging (FUNCAT – decrease in FUNCAT signal implies translation inhibition) (**Fig 1A**) ^43–45^. In the first experiment, we treated the rat primary cortical neurons with neurobasal (control) or conditioned neurobasal media from APOE KO or APOE3 or APOE4 iPSCs (final concentration of APOE was 10-15nM) for 20 minutes (**Fig 1B**). APOE4 conditioned media treatment for 20 minutes caused a significant 1.5-fold increase in eEF2 phosphorylation compared to APOE3/ APOE KO/ untreated conditions (**Fig 1B**). Treatment with APOE3 (the most common APOE variant in the human population) did not cause any change in eEF2 phosphorylation compared to APOE KO treatment or untreated conditions (**Fig 1B**). This indicated that there could be a significant reduction of protein synthesis in the neurons exposed to APOE4 for 20 minutes. The same response was observed when the neurons were treated with recombinant APOE protein instead of conditioned media at a similar concentration (15nM). With recombinant APOE4 treatment for 20 minutes, a significant 2-fold increase in eEF2 phosphorylation was observed compared to both untreated and APOE3 treated neurons (**Fig 1C**). Since the phosphorylation of eEF2 indicates the translation status at a given time point, we used the FUNCAT assay to measure the cumulative effect of APOE4 exposure on protein synthesis for 20 minutes in neurons. The FUNCAT assay was validated in neurons that were exposed to methionine analog AHA but without the follow-up click chemistry (**Fig S1H**). Neurons treated with recombinant APOE4 protein for 20 minutes showed a significantly reduced FUNCAT signal compared to APOE3 treated or untreated neurons (**Fig 1D and 1E**) indicating a decreased protein synthesis in neurons exposed to APOE4. We also observed an increase in ERK phosphorylation in neurons exposed to APOE4 conditioned media or APOE4 recombinant protein for 20 minutes (**Fig S1F and S1G**), a feature reported previously^37,38^, thus validating our APOE treatment paradigm.

**Figure 1.**
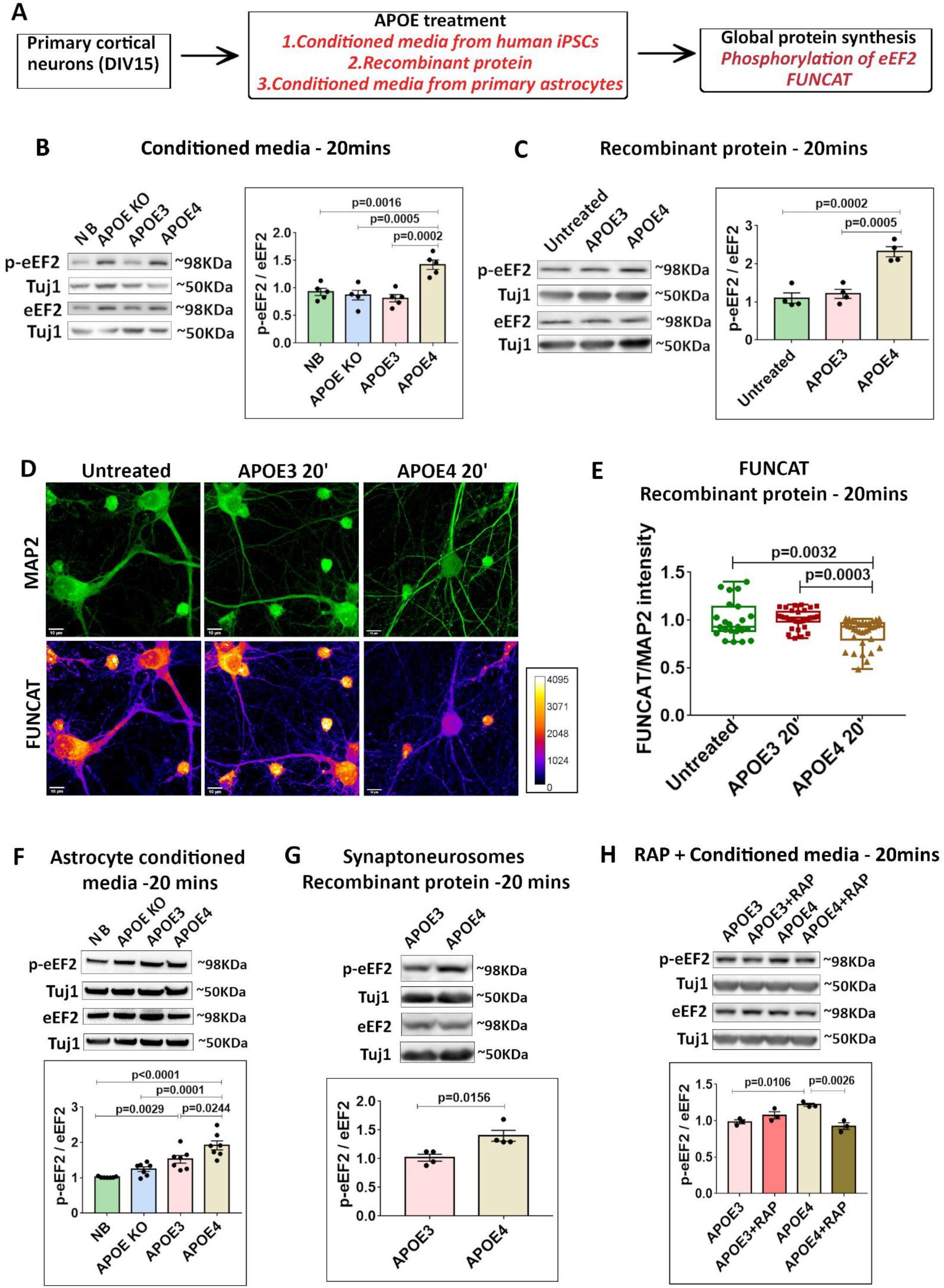
APOE4 treatment for 20 minutes decreases global protein synthesis in neurons and synpatoneurosomes. **A** – Experimental workflow. Primary rat/mouse cortical neurons (DIV15) were treated with APOE from different sources - stem cell secreted APOE, astrocyte secreted APOE and recombinant APOE protein. The APOE treated neurons were probed for readouts of global protein synthesis. **B** - Rat primary cortical neurons (DIV15) were treated with APOE (10-15nM) from iPSC conditioned media for 20 minutes and probed for phosphorylation of eEF2. Left - representative immunoblots indicating levels of phospho-eEF2, eEF2 and Tuj1; Right - graph indicating ratio of phospho-eEF2 to eEF2 normalized to Tuj1. Data is represented as mean +/- SEM. N=5, One-way ANOVA (p=0.0001) followed by Tukey’s multiple comparison test. **C** - Rat primary cortical neurons (DIV15) were treated with recombinant APOE protein (15nM) for 20 minutes and probed for phosphorylation of eEF2. Left - representative immunoblots indicating levels of phospho-eEF2, eEF2 and Tuj1; Right - graph indicating ratio of phospho-eEF2 to eEF2 normalized to Tuj1. Data is represented as mean +/- SEM. N=4, One-way ANOVA (p=0.0002) followed by Tukey’s multiple comparison test. **D** – Rat primary cortical neurons (DIV15) were treated with recombinant APOE protein (15nM) for 20 minutes and subjected to fluorescent non-canonical amino acid tagging (FUNCAT) along with immunostaining for MAP2. The representative images for MAP2 and FUNCAT fluorescent signal under different APOE treatment conditions are shown (Scale bar - 10μM). **E** – The graph represents the quantification of the FUNCAT fluorescent intensity normalized to MAP2 fluorescent intensity under different APOE treatment conditions. Data is represented as mean +/- SEM. N = 20-40 neurons from 4 independent experiments, One-way ANOVA (p=0.0001) followed by Tukey’s multiple comparison test. **F** - Mouse primary cortical neurons (DIV15) were treated with APOE from mouse primary astrocyte conditioned media for 20 minutes and probed for phosphorylation of eEF2. Top - representative immunoblots indicating levels of phospho-eEF2, eEF2 and Tuj1; Bottom - graph indicating ratio of phospho-eEF2 to eEF2 normalized to Tuj1. Data is represented as mean +/- SEM. N=7, One-way ANOVA (p<0.0001) followed by Tukey’s multiple comparison test. **G** – Synaptoneurosomes prepared from P30 rat cortices were treated with recombinant APOE protein (15nM) for 20 minutes and probed for phosphorylation of eEF2. Top - representative immunoblots indicating levels of phospho-eEF2, eEF2 and Tuj1; Bottom - graph indicating ratio of phospho-eEF2 to eEF2 normalized to Tuj1. Data is represented as mean +/- SEM. N=4, Unpaired Student’s t-test. **H** - Rat primary cortical neurons (DIV15) were treated with APOE receptor antagonist RAP (200nM) along with APOE (10-15nM) from iPSC conditioned media for 20 minutes and probed for phosphorylation of eEF2. Top - representative immunoblots indicating levels of phospho-eEF2, eEF2 and Tuj1; Bottom - graph indicating ratio of phospho-eEF2 to eEF2 normalized to Tuj1. Data is represented as mean +/- SEM. N=5, One-way ANOVA (p=0.003) followed by Tukey’s multiple comparison test.

Since APOE is primarily secreted by astrocytes in the brain, we wanted to investigate if the APOE4 secreted from in-vitro cultured astrocytes has the same effect on neuronal translation as the APOE4 secreted by iPSCs. To study this, we cultured primary astrocytes from APOE KO mice or humanized APOE 3/3 knock-in mice or humanized APOE 4/4 knock-in mice. On DIV12 of the astrocyte culture, the astrocyte media was replaced by neurobasal and the conditioned media was collected after 48 hours. The primary cultured neurons (DIV 15) from C57Bl/6 mice were exposed to the mouse APOE astrocyte conditioned media for 20 minutes and the translation status was measured by the phosphorylation of eEF2. We observed a significant increase in the phosphorylation of eEF2 in mouse neurons exposed to APOE4 astrocyte conditioned media as compared to APOE3 astrocyte conditioned media / APOE KO astrocyte conditioned media / neurobasal conditions (**Fig 1F**), further validating the effect of APOE4 on translation. To test whether the APOE4 induced translation inhibition occurs in the synaptic compartments, we prepared synaptoneurosomes from P30 rat cortices and treated them with 15nM recombinant APOE protein for 20 minutes. We observed a significant increase in the phosphorylation of eEF2 (**Fig 1G**) and ERK (**Fig S1K**) in the synaptoneurosomes exposed to APOE4 recombinant protein as compared to APOE3, corroborating that the APOE4 mediated effect was synaptic as well. Lastly, we verified our findings in the human neuron system. APOE KO iPSCs were differentiated into forebrain glutamatergic APOE KO neurons using monolayer dual-SMAD inhibition protocol^46,47^. Human APOE KO neurons (1 month into neuronal maturation) were treated with APOE3 or APOE4 iPSC conditioned media for 20 minutes and probed for eEF2 phosphorylation (**Fig S1I**). APOE4 treatment of 20 minutes caused a significant increase in eEF2 phosphorylation compared to APOE3 treatment in the human neurons as well (**Fig S1J**), further supporting the finding of global translation inhibition by APOE4.

Finally, to confirm that this effect on translation is due to the binding of APOE to its cognate receptor, we inhibited the APOE receptor using RAP (Receptor Associated Protein). In the presence of RAP, APOE4 induced increase in the phosphorylation of eEF2 was absent (**Fig 1H**). Similarly, RAP was also able to completely prevent the APOE4 mediated increase in the phosphorylation of ERK (**Fig S1L**). In summary, we have shown that APOE4 exposure for 20 minutes caused a significant reduction of protein synthesis in neurons and synaptoneurosomal preparations. We have demonstrated this in different systems and by using APOE from multiple sources including iPSC conditioned media, astrocyte conditioned media, and recombinant protein. We have also validated that the translation inhibition is specifically due to the binding of APOE to its cognate receptors in the neurons.

### NMDAR mediated translation response is completely lost on APOE4 exposure

APOE4 was previously reported to interfere with NMDAR signalling^39,41,48^ and we observed a significant inhibition of translation in neurons exposed to APOE4 (**Fig 1**). Hence, we aimed to investigate if the NMDAR mediated protein synthesis response is affected by the presence of APOE4. The activity of NMDA receptors has a very important role in synaptic plasticity. NMDAR stimulation leads to a distinct translation response which initially involves an inhibition of global translation with a small subset of mRNAs being translationally upregulated and a more robust translation activation at a later phase^44,49,50^. Stimulation of cortical neurons with 20μM NMDA for 5 minutes led to an increase in phosphorylation of eEF2 as previously reported^44,49^ (**Fig S2A**), indicating global translation inhibition. In this background, the translation of specific candidates such as PTEN and PSD-95 is known to get activated^49^. Consequently, we observed an increase in the protein levels of PTEN (**Fig S2C**) and PSD-95 (**Fig S2E**) with our stimulation paradigm in the neurons, clearly representing NMDAR specific translation response.

To test the effect of APOE treatment on NMDAR mediated translation response, we incubated rat cortical neurons with APOE KO/ APOE3/ APOE4 iPSC conditioned media (for biochemistry experiments) or recombinant protein (for FUNCAT experiment) for 20 minutes as previously described. During the last 5 minutes, the neurons were incubated with 20μM NMDA, followed by different assays to study the translation response (**Fig 2A**). When we incubated the neurons with APOE KO conditioned media for 20 minutes, the translation response to NMDAR stimulation was unaffected. We observed an increase in eEF2 phosphorylation (**Fig S2B**) and an increase in PTEN (**Fig S2D**) and PSD-95 (**Fig S2F**) protein levels. NMDAR stimulation of the APOE3 conditioned media treated neurons showed a similar significant increase in the phosphorylation of eEF2 indicating a normal NMDAR response (**Fig 2B**). On the other hand, the neurons exposed to APOE4 conditioned media did not show an increase in eEF2 phosphorylation on NMDAR stimulation (**Fig 2B**). It should be noted that APOE4 treatment alone caused an increase in eEF2 phosphorylation compared to APOE3 treated neurons, but APOE4 treated neurons failed to evoke NMDAR mediated increase in eEF2 phosphorylation (**Fig 2B**). We have previously shown that NMDAR stimulation for 5 minutes causes a significant decrease in the total protein synthesis as measured by FUNCAT assay^44^. In the neurons treated with recombinant APOE3 protein, 5-minute stimulation of NMDA receptors led to a decrease in the FUNCAT signal (**Fig 2C and 2D**) indicating a normal NMDAR response. But, in the neurons exposed to recombinant APOE4 protein, the NMDA induced decrease in the FUNCAT signal was completely lost (**Fig 2C and 2D**). Again, APOE4 exposure alone caused a significant decrease in the FUNCAT signal and NMDAR stimulation failed to elicit a response in this background.

**Figure 2.**
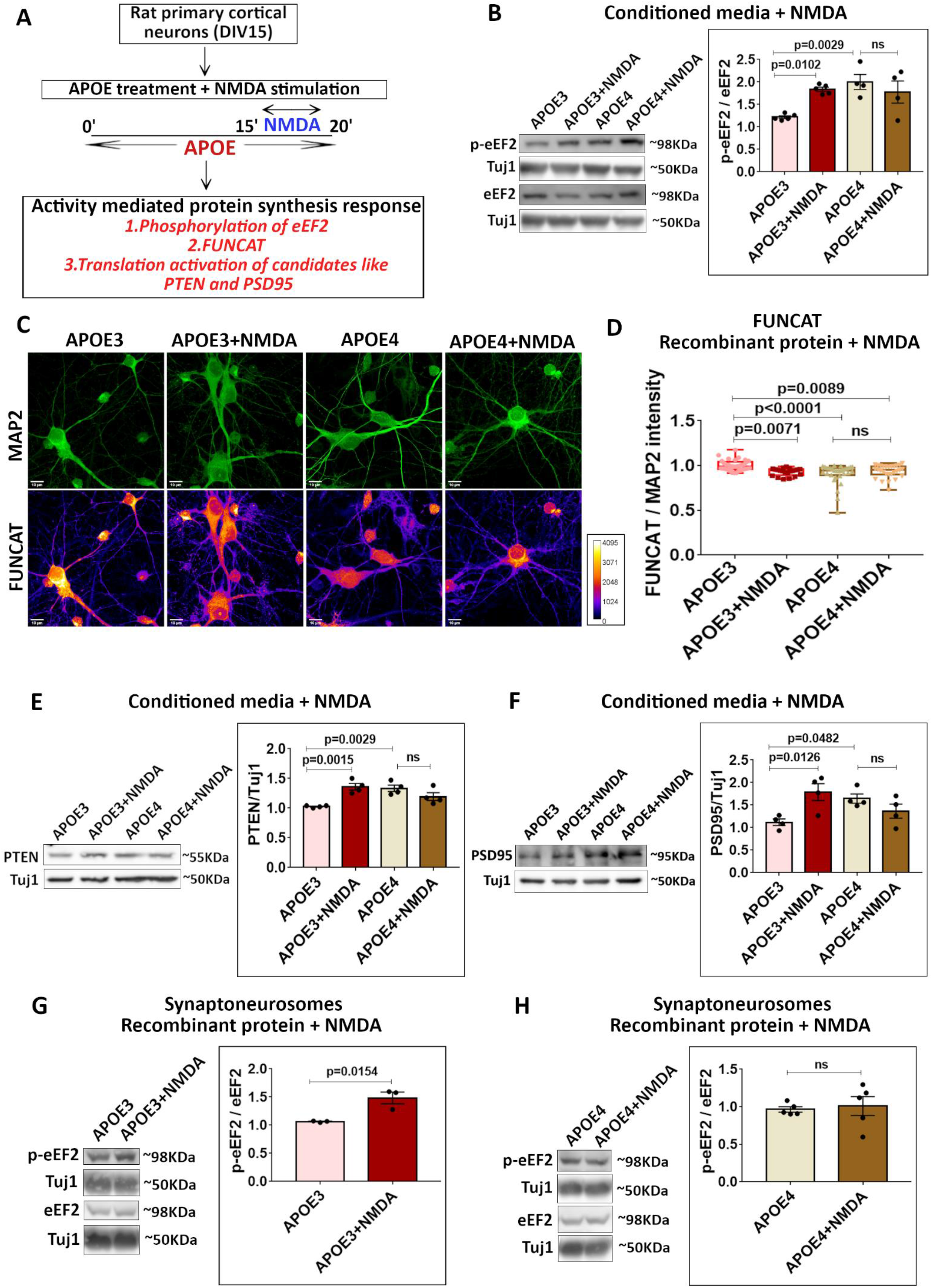
The NMDAR mediated translation response is lost in APOE4 treated neurons. **A** – Experimental workflow. Rat primary cortical neurons (DIV15) were treated with APOE from iPSC conditioned media/recombinant protein for 20 minutes. During the last 5 minutes of the APOE treatment, the neurons were subjected to stimulation of NMDA receptors (20μM NMDA, 5 minutes). The NMDAR-mediated translation response in APOE treated neurons was probed by using the following readouts - phosphorylation eEF2, FUNCAT and increased levels of PTEN and PSD95 proteins. **B** - Rat primary cortical neurons (DIV15) were treated with APOE3/APOE4 (10-15nM) conditioned media for 20 minutes along with NMDAR stimulation for 5 minutes (20μM NMDA). The cell lysates were probed for phosphorylation of eEF2. Left - representative immunoblots indicating levels of phospho-eEF2, eEF2 and Tuj1; Right - graph indicating ratio of phospho-eEF2 to eEF2 normalized to Tuj1. Data is represented as mean +/- SEM. N=4-5, One-way ANOVA (p=0.0047) followed by Dunnett’s multiple comparison test. **C** - Rat primary cortical neurons (DIV15) were treated with recombinant APOE3 or APOE4 protein (15nM) for 20 minutes along with NMDAR stimulation for 5 minutes (20μM NMDA). They were subjected to fluorescent non-canonical amino acid tagging (FUNCAT) along with immunostaining for MAP2. The representative images for MAP2 and FUNCAT fluorescent signal under different treatment conditions are shown (Scale bar - 10μM). **D** – The graph represents the quantification of the FUNCAT fluorescent intensity normalized to MAP2 fluorescent intensity under different treatment conditions. Data is represented as mean +/- SEM. N = 20-40 neurons from 2 independent experiments, One-way ANOVA (p<0.0001) followed by Tukey’s multiple comparison test. **E** - Rat primary cortical neurons (DIV15) were treated with APOE3/APOE4 (10-15nM) conditioned media for 20 minutes along with NMDAR stimulation for 5 minutes (20μM NMDA). The cell lysates were probed for PTEN protein. Left - representative immunoblots indicating levels of PTEN and Tuj1; Right - graph indicating PTEN levels normalized to Tuj1. Data is represented as mean +/- SEM. N=4, One-way ANOVA (p=0.002) followed by Dunnett’s multiple comparison test. **F** - Rat primary cortical neurons (DIV15) were treated with APOE3/APOE4 (10-15nM) conditioned media for 20 minutes along with NMDAR stimulation for 5 minutes (20μM NMDA). The cell lysates were probed for PSD95 protein. Left - representative immunoblots indicating levels of PSD95 and Tuj1; Right - graph indicating PSD95 levels normalized to Tuj1. Data is represented as mean +/- SEM. N=4, One-way ANOVA (p=0.0216) followed by Dunnett’s multiple comparison test. **G** - Synaptoneurosomes prepared from P30 rat cortices were treated with recombinant APOE3 protein (15nM) for 20 minutes along with NMDAR stimulation for 5 minutes (40μM NMDA). The lysates were probed for phosphorylation of eEF2. Left - representative immunoblots indicating levels of phospho-eEF2, eEF2 and Tuj1; Right - graph indicating ratio of phospho-eEF2 to eEF2 normalized to Tuj1. Data is represented as mean +/- SEM. N=3, Unpaired Student’s t-test. **H** - Synaptoneurosomes prepared from P30 rat cortices were treated with recombinant APOE4 protein (15nM) for 20 minutes along with NMDAR stimulation for 5 minutes (40μM NMDA). The lysates were probed for phosphorylation of eEF2. Left - representative immunoblots indicating levels of phospho-eEF2, eEF2 and Tuj1; Right - graph indicating ratio of phospho-eEF2 to eEF2 normalized to Tuj1. Data is represented as mean +/- SEM. N=5, Unpaired Student’s t-test.

Regarding the NMDAR specific translation activation, APOE3 conditioned media treated neurons showed a significant increase in PTEN (**Fig 2E**) and PSD-95 (**Fig 2F**) protein levels on 5-minute NMDAR stimulation. But, in the presence of APOE4 conditioned media, NMDAR mediated increase of PTEN and PSD-95 proteins were lost (**Fig 2E and 2F**). Furthermore, there were elevated levels of PTEN and PSD-95 proteins on APOE4 exposure alone but NMDAR mediated increase was not observed. Since APOE4 has been previously reported to lead to transcriptional activation of several synaptic mRNAs^38^, we checked whether PTEN and PSD-95 mRNA levels were altered by APOE exposure. We did not observe any change in the mRNA levels of both PTEN (**Fig S2G**) and PSD-95 (**Fig S2H**) on APOE KO/ APOE3/ APOE4 iPSC conditioned media treatment.

Finally, we also tested NMDAR response in the presence of APOE in rat cortical synaptoneurosome preparations. When synaptoneurosomes were exposed to recombinant APOE3 protein for 20 minutes, they responded to NMDAR stimulation by showing an increase in eEF2 phosphorylation (**Fig 2G**). But 20-minute exposure to APOE4 recombinant protein resulted in a complete loss of the NMDAR response as there was no change in the phosphorylation of eEF2 (**Fig 2H**). In summary, 20 minutes of exposure to APOE4 completely abolished NMDAR mediated translation response with respect to both global translation inhibition and translation activation of specific candidates, whereas NMDAR mediated translation response is unaffected with 20-minute APOE3 treatment. Interestingly, 20-minute treatment with APOE4 seemed to elicit the 5-minute NMDA stimulation mediated translation response implying a role of NMDARs in the APOE4 mediated synaptic defect.

### Aberrant translation response by APOE4 mimics the translation response upon 5-minute NMDAR stimulation

It was interesting that whilst 20 minutes of APOE4 exposure abolished the NMDAR translation response in neurons, APOE4 treatment alone mimicked NMDAR translation response (**Fig 2**). To further investigate this observation, we used the polysome profiling assay. We separated the rat primary neuron lysate on a 15 - 45% linear sucrose gradient which was fractionated into 11 samples (**Fig 3A**). We used puromycin treatment to identify the translationally active polysome fractions (**Fig 3B, S3A, and S3B**). Puromycin treatment disrupted the ribosomal fractions 7-11 (as indicated by the immunoblot for ribosomal protein RPLP0) indicating that they are the actively translating pool (**Fig 3B, S3A and S3B**). Fractions 1-6 which are insensitive to puromycin treatment were grouped as non-translating pool (**Fig 3B, S3A and S3B**). On 5-minute stimulation with 20μM NMDA, we observed an increase in eEF2 phosphorylation indicating global translation inhibition (**Fig 2**). This global translation inhibition was also clearly reflected in the shifting of ribosomes (RPLP0 protein) from the translationally active pool (fractions 7-11) to the inactive pool (fractions 1-6) (**Fig 3C and S3C)**. 20-minute treatment with APOE3 conditioned media along with 5-minute NMDAR stimulation showed the same response of ribosomal shift (RPLP0) towards the translationally inactive fractions (**Fig 3D, 3E, and S3D**), indicating that exposure to APOE3 does not affect the NMDAR response. APOE4 conditioned media exposure for 20 minutes resulted in the same shift of ribosomes (RPLP0) into the translationally inactive pool (**Fig 3D, 3E, and S3D**), similar to 5-minute NMDAR stimulation in the APOE3 background. Interestingly, APOE4 treatment for 20 minutes along with 5-minute NMDAR stimulation caused a further and maximum shift of the ribosomes (RPLP0) from the translationally active fractions 7-11 towards the inactive fractions 1-6 (**Fig 3D, 3E, and S3D**).

**Figure 3.**
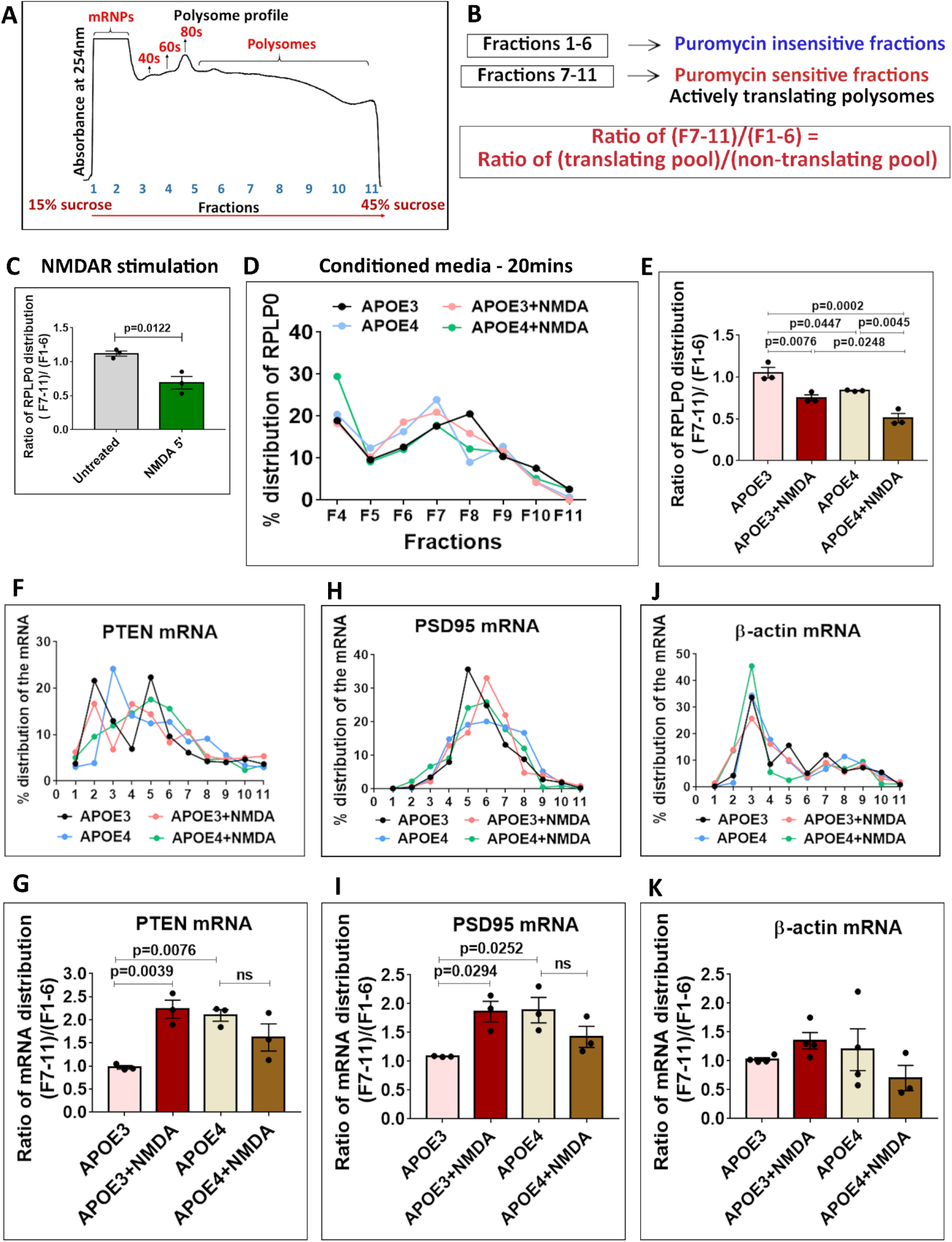
Treatment of neurons with APOE4 mimics the 5-minute NMDAR response of global protein synthesis inhibition and translation activation of specific candidate mRNAs. **A** – Representative polysome profile or absorbance profile at 254nm indicating the mRNPs, ribosomal subunits (40S,60S), monosomes (80S) and polysomes distributed on 15% to 45% linear sucrose gradient. **B** – Schematic showing the division of polysome profiling fractions into two pools based on sensitivity to puromycin treatment. Fractions 7-11 was the puromycin sensitive or actively translating pool and Fractions 1-6 was the puromycin insensitive pool or non-translating pool. The ratio of the percentage of protein or mRNA in Fractions 7-11/Fractions 1-6 indicates their distribution in the translating pool/non-translating pool. **C** - Rat primary cortical neurons (DIV15) were subjected to NMDAR stimulation for 5 minutes (20μM NMDA). The cell lysates were subjected to polysome profiling and probed for the distribution of ribosomal protein RPLP0. The graph represents the ratio of RPLP0 protein distribution in fractions 7-11 (translating pool) to fractions 1-6 (non-translating pool). Data is represented as mean +/- SEM. N=3, Unpaired Student’s t-test. **D** - Rat primary cortical neurons (DIV15) were treated with APOE3/APOE4 (10-15nM) conditioned media for 20 minutes along with NMDAR stimulation for 5 minutes (20μM NMDA). The cell lysates were subjected to polysome profiling and probed for the distribution of ribosomal protein RPLP0 across the linear sucrose gradient. The line graph represents the percentage distribution of RPLP0 protein in Fractions 4-11 under different treatment conditions. **E** – The graph represents the ratio of RPLP0 protein distribution in fractions 7-11 (translating pool) to fractions 1-6 (non-translating pool) under different APOE treatment conditions. Data is represented as mean +/- SEM. N=3, One-way ANOVA (p=0.0002) followed by Tukey’s multiple comparison test. Rat primary cortical neurons (DIV15) were treated with APOE3/APOE4 (10-15nM) conditioned media for 20 minutes along with NMDAR stimulation for 5 minutes (20μM NMDA). The cell lysates were subjected to polysome profiling and probed for the distribution of the following candidate mRNAs in each fraction using RT-PCR method – **F** - The line graph represents the percentage distribution of PTEN mRNA in each fraction under different treatment conditions. **G** - The graph represents the ratio of PTEN mRNA distribution in fractions 7-11 (translating pool) to fractions 1-6 (non-translating pool). Data is represented as mean +/- SEM. N=3, One-way ANOVA (p=0.006) followed by Dunnett’s multiple comparison test. **H** - The line graph represents the percentage distribution of PSD95 mRNA in each fraction under different treatment conditions. **I** - The graph represents the ratio of PSD95 mRNA distribution in fractions 7-11 (translating pool) to fractions 1-6 (non-translating pool). Data is represented as mean +/- SEM. N=3, One-way ANOVA (p=0.0286) followed by Dunnett’s multiple comparison test. **J** - The line graph represents the percentage distribution of β-actin mRNA in each fraction under different treatment conditions. **K** - The graph represents the ratio of β-actin mRNA distribution in fractions 7-11 (translating pool) to fractions 1-6 (non-translating pool). Data is represented as mean +/- SEM. N=3, One-way ANOVA (ns).

We have previously shown that, in the background of global inhibition of protein synthesis, NMDAR stimulation leads to translation activation of specific candidates such as PTEN and PSD-95 (**Fig 2E and 2F**)^49^. We further investigated the distribution of PTEN and PSD-95 mRNAs on the polysome profile in the presence of APOE. On 5-minute NMDAR stimulation in the APOE3 background (20 minutes), there was a shift of PTEN and PSD-95 mRNAs into translationally active polysome fractions (**Fig 3F, 3G, and 3H, 3I**) validating that the normal NMDAR response was intact in the background of APOE3. Conversely, PTEN and PSD-95 mRNAs shifted to the translating pool with 20 minutes of APOE4 exposure, even without NMDAR stimulation (**Fig 3F, 3G and 3H, 3I**). This reinforces the idea that 20 minutes of exposure to APOE4 brings about the same translation response as 5-minute NMDAR stimulation from the perspectives of both global translation inhibition and translation activation of specific candidates. Additionally, NMDAR stimulation in the background of APOE4 elicited a very different response. It did not lead to the translation activation of PTEN and PSD-95 mRNAs (**Fig 3F, 3G, and 3H, 3I**), though it caused a further decrease of ribosomes from the translationally active pool compared to APOE4 treatment (**Fig 3D and 3E**). β-actin and α-tubulin mRNAs were used as controls as their translation is unaffected by both NMDAR stimulation and APOE exposure (**Fig 3J, 3K and S3E, S3F**). In summary, these results indicated that 20-minute treatment with APOE4 causes a translation response which appeared to be similar to 5-minute NMDAR stimulation under control or APOE3 conditions. However, NMDAR stimulation in the background of APOE4 resulted in an aberrant translation response.

### NMDAR stimulation in the background of APOE4 causes a stress response

Following 20-minute treatment with APOE4, neurons lost the ability to respond to NMDAR stimulation whilst similar exposure to APOE3 had no such effect (**Fig 2 and 3**). In the polysome profiling assay, we observed that the shift of ribosomes into the translationally inactive pool was highest in the neurons stimulated with NMDA in the background of APOE4 (**Fig 3D and 3E)**. To understand this phenomenon better, we further analyzed only the puromycin insensitive fractions (Fractions 1-6) to study the redistribution of the mRNAs within this pool. Within this pool, Fractions 4-6 showed a signal for the ribosomal proteins RPLP0 (large subunit) or RPS6 (small subunit) or both indicating that these fractions contain either ribosomal subunits or monosomes, as reflected in the A254 trace as well (**Fig 4A and 4B**). Fractions 1-3, which did not show any signal for ribosomal proteins, were mainly enriched in the mRNPs (messenger ribonucleoproteins) representing the mRNAs that are not associated with the ribosomes.

**Figure 4.**
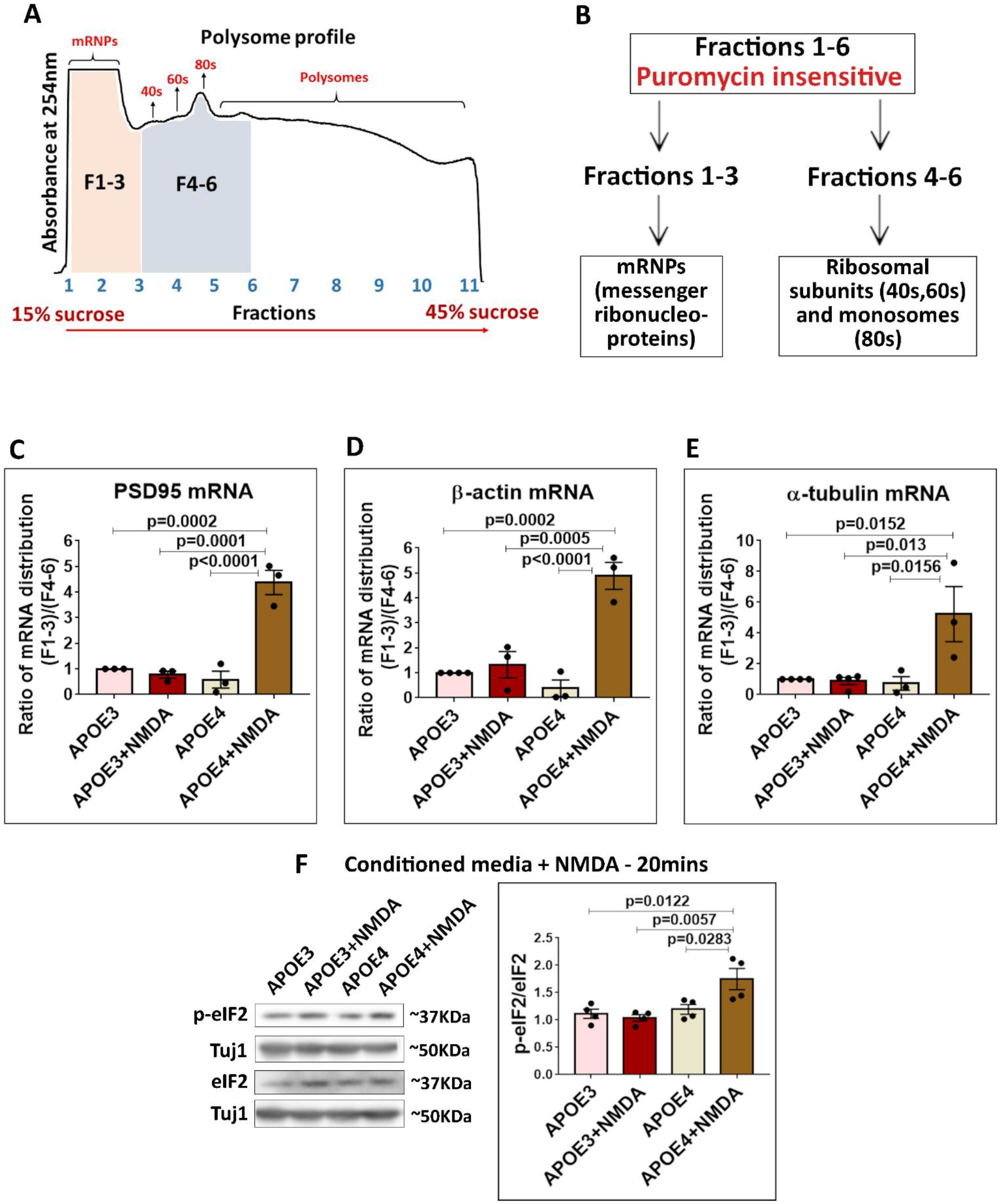
The NMDAR stimulation of APOE4 treated neurons invokes a stress response, potentially through inhibition of translation initiation and eIF2 phosphorylation. **A** – Polysome profile indicating the fractions (F1-3 and F4-6) from the puromycin insensitive pool considered for the experiments and analysis. **B** – Schematic showing division of puromycin insensitive fractions 1-6 into two pools -fractions 1-3 which constitute the mRNPs and fractions 4-6 which constitute the ribosomal subunits (40s,60s) and monosomes (80s). Rat primary cortical neurons (DIV15) were treated with APOE3/APOE4 (10-15nM) conditioned media for 20 minutes along with NMDAR stimulation for 5 minutes (20μM NMDA). The cell lysates were subjected to polysome profiling and probed for the distribution of the following candidate mRNAs in fractions 1-6 using RT-PCR method – **C** - The graph represents the ratio of PSD95 mRNA distribution in fractions 1-3 to fractions 4-6. Data is represented as mean +/- SEM. N=3, One-way ANOVA (p<0.0001) followed by Tukey’s multiple comparison test. **D** - The graph represents the ratio of β-actin mRNA distribution in fractions 1-3 to fractions 4-6. Data is represented as mean +/- SEM. N=3, One-way ANOVA (p<0.0001) followed by Tukey’s multiple comparison test. **E** - The graph represents the ratio of α-tubulin mRNA distribution in fractions 1-3 to fractions 4-6. Data is represented as mean +/- SEM. N=3-4, One-way ANOVA (p=0.0079) followed by Tukey’s multiple comparison test. **F** - Rat primary cortical neurons (DIV15) were treated with APOE3/APOE4 (10-15nM) conditioned media for 20 minutes along with NMDAR stimulation for 5 minutes (20μM NMDA) and probed for phosphorylation of eIF2α. Left - representative immunoblots indicating levels of phospho-eIF2α, eIF2α and Tuj1; Right - graph indicating ratio of phospho-eIF2 to eIF2 normalized to Tuj1. Data is represented as mean +/- SEM. N=4, One-way ANOVA (p=0.0047) followed by Tukey’s multiple comparison test.

When we investigated the redistribution of the mRNAs between mRNP fractions (fractions 1-3) and ribosomal fractions (fractions 4-6) on NMDAR stimulation, we did not observe any significant change in the APOE3 background, whilst there was a huge increase of mRNAs in the mRNP fractions (fractions 1-3) in the APOE4 background (**Fig 4C, 4D and 4E**). While NMDAR mediated translation activation was specific only to a subset of mRNAs, there was no such specificity in the NMDA induced shift of mRNAs into the mRNP fractions in the APOE4 background. Even the mRNAs such as β-actin and α-tubulin which did not respond to NMDAR stimulation in the APOE3 background were shifted to mRNP fractions in the APOE4 background (**Fig 4D and 4E**). This was an aberrant response that was consistent with stress-induced global translation inhibition.

To verify if this a was stress response, we probed into the phosphorylation status of eukaryotic translation initiation factor eIF2α which is a well-established marker for stress^51^. APOE3 conditioned media treatment (20 minutes) along with 5-minute NMDAR stimulation did not cause a change in the phosphorylation of eIF2α (**Fig 4F**) while it increased eEF2 phosphorylation (**Fig 2**). But, in the presence of APOE4 conditioned media (20 minutes), 5-minute NMDAR stimulation caused a significant increase in eIF2α phosphorylation (**Fig 4F**) while there was no change in eEF2 phosphorylation (**Fig 2**). Additionally, NMDAR stimulation in the background of APOE4 caused a clear shift of ribosomal protein RPS6 (small subunit) into fractions 3-4 (**Fig S4A, S4B, and S4C**) which very likely represents the unassembled subunits of the ribosome. The accumulation of the small subunit of the ribosome indicated an increased inhibition of translation at the initiation step, which was another characteristic feature of the stress response. We did not observe such a shift of RPS6 on NMDAR stimulation in the APOE3 background (**Fig S4A, S4B, and S4C**). Overall, these results indicated that NMDAR stimulation of APOE4 treated neurons potentially leads to a pathological stress response.

### Translation response elicited by APOE treatment and NMDAR stimulation are different temporally

An important feature of synaptic activity mediated translation is its unique spatio-temporal pattern^44^. NMDAR mediated translation response is a good example of this. Initially, NMDAR stimulation leads to a global translation inhibition which over time turns to a robust translation activation phase^44^. We were able to capture this temporal pattern of NMDAR mediated translation response using the phosphorylation status of eEF2 as a readout (**Fig 5A**). On NMDAR stimulation (20μM) of cultured cortical neurons, there was a rapid and robust 2-fold increase in the phosphorylation of eEF2 at 1 minute. By 5 minutes, the phosphorylation of eEF2 was reduced, but it remained significantly high (1.5-fold increase) compared to the untreated/basal condition. By 20 minutes, the phosphorylation of eEF2 was further reduced below the basal/untreated level (>1.0) indicating global translation activation (**Fig 5A**). This temporal pattern of translation regulation was also reflected in the polysome profiling assay and RPLP0 distribution (**Fig 5B and S5A**). Stimulation with NMDA (20μM) for 1 minute did not cause a significant change in the shift of ribosomes (RPLP0 distribution). However, 20-minute stimulation with NMDA (20μM) caused a significant increase in the heavier polysomes (RPLP0 shift towards translating fractions) clearly indicating the phase of translation activation (**Fig 5B and S5A**).

**Figure 5.**
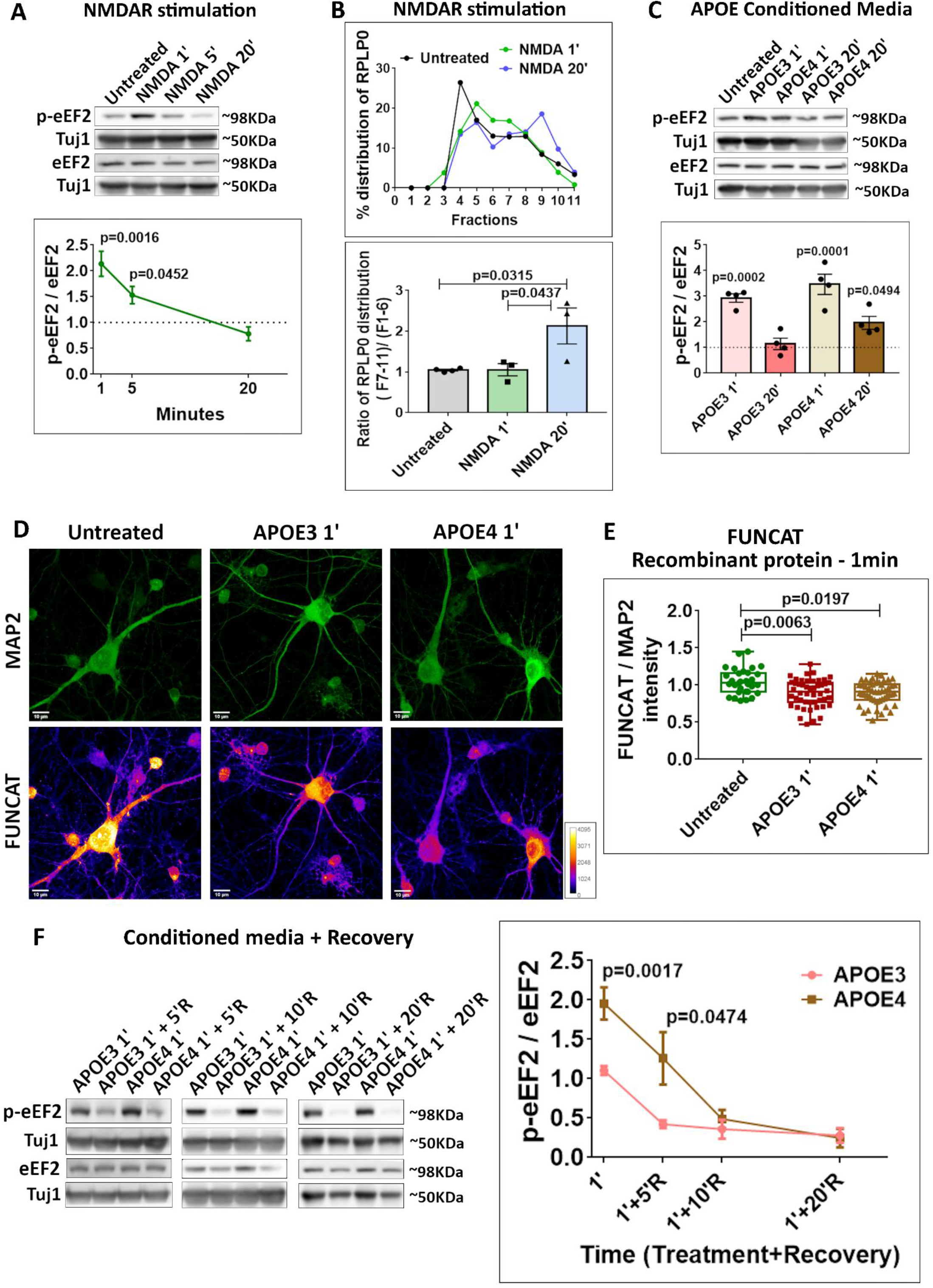
Temporal profiles and recovery kinetics of APOE3 and APOE4 translation response. **A** - Rat primary cortical neurons (DIV15) were treated with NMDA (20μM) for 1 minute, 5 minutes, 20 minutes and probed for phosphorylation of eEF2. Top - representative immunoblots indicating levels of phospho-eEF2, eEF2 and Tuj1; Bottom - graph indicating ratio of phospho-eEF2 to eEF2 normalized to Tuj1. Data is represented as mean +/- SEM. N=3, One-way ANOVA (p=0.0015) followed by Tukey’s multiple comparison test. **B** - Rat primary cortical neurons (DIV15) were subjected to NMDAR stimulation for 1 minute and 20 minutes (20μM NMDA). The cell lysates were subjected to polysome profiling and probed for the distribution of ribosomal protein RPLP0. Top – line graph showing the percentage distribution of RPLP0 under different conditions. Bottom - bar graph representing the ratio of RPLP0 protein distribution in fractions 7-11 (translating pool) to fractions 1-6 (non-translating pool). Data is represented as mean +/- SEM. N=3. One-way ANOVA (p=0.0248) followed by Tukey’s multiple comparison test. **C** - Rat primary cortical neurons (DIV15) were treated with APOE3/APOE4 (10-15nM) conditioned media for 1 minute and 20 minutes and probed for phosphorylation of eEF2. Top - representative immunoblots indicating levels of phospho-eEF2, eEF2 and Tuj1; Bottom - graph indicating ratio of phospho-eEF2 to eEF2 normalized to Tuj1. Data is represented as mean +/- SEM. Dotted line indicates the untreated condition. All the APOE treatments were normalized to untreated condition. N=4. One-way ANOVA (p<0.0001) followed by Dunnett’s multiple comparison test. **D** – Rat primary cortical neurons (DIV15) were treated with recombinant APOE protein (15nM) for 1 minute and subjected to fluorescent non-canonical amino acid tagging (FUNCAT) along with immunostaining for MAP2. The representative images for MAP2 and FUNCAT fluorescent signal under different APOE treatment conditions are shown (Scale bar - 10μM). **E** – The graph represents the quantification of the FUNCAT fluorescent intensity normalized to MAP2 fluorescent intensity under different APOE treatment conditions. Data is represented as mean +/- SEM. N = 20-40 neurons from 4 independent experiments, One-way ANOVA (p=0.0058) followed by Tukey’s multiple comparison test. **F** - Rat primary cortical neurons (DIV15) were treated with APOE3/APOE4 (10-15nM) conditioned media for 1 minute and subjected to recovery for 5 minutes/ 10 minutes / 20 minutes using pre-conditioned neurobasal media. The samples were probed for the phosphorylation of eEF2. The data from each recovery time point is normalized to its corresponding 1-minute APOE treated set. Left - representative immunoblots indicating levels of phospho-eEF2, eEF2 and Tuj1; Right - graph indicating ratio of phospho-eEF2 to eEF2 normalized to Tuj1. Data is represented as mean +/- SEM. For APOE3 1’ and APOE4 1’, N=8, Unpaired Student’s t-test. For APOE 1’ + 5’R, N=4, Unpaired Student’s t-test. For APOE 1’ + 10’R N=5. For APOE 1’ + 20’R, N=3.

In the previous sections, we have shown that exposure to APOE4 (but not APOE3) for 20 minutes generated a translation response similar to 5-minute NMDAR stimulation in two important aspects - global translation inhibition and translation activation of specific candidates. For a more comprehensive understanding of the effect of APOE in comparison to NMDAR stimulation, we studied the effect of APOE3 and APOE4 on neuronal translation at both 1-minute and 20-minute time points. Interestingly, with 1-minute conditioned media exposure, unlike the 20-minute time point, both APOE3 and APOE4 showed a significant increase in the eEF2 phosphorylation (**Fig 5C**). There was also a significant decrease in the FUNCAT signal in neurons exposed to recombinant APOE3 and APOE4 for 1 minute (**Fig 5D and 5E**). These results indicated that at the 1-minute time point, both APOE3 and APOE4 elicited a translation response that resembled NMDAR stimulation. The key difference was at the 20-minute time point where the phosphorylation of eEF2 was below the basal level (~1.35-fold) on NMDAR stimulation (**Fig 5A**) indicating translation activation. In the case of APOE3, the phosphorylation of eEF2 was back to basal level by 20 minutes (**Fig 5C**). With APOE4 exposure, though eEF2 phosphorylation at 20-minute was lower than the 1-minute time point, it was still significantly higher (1.5-fold) than the basal level (**Fig 5C**). This indicated that the recovery from the translation inhibition could be different between APOE3 and APOE4 treatments. Unlike the 20-minute time point where APOE4 treated neurons showed a shift of ribosomes to the non-translating fractions compared to APOE3 treatment (**Fig 3**), there was no difference in the ribosome (RPLP0) distribution between APOE3 and APOE4 conditioned media treated neurons at 1 minute (**Fig S5E**).

Since the brain has elaborate mechanisms to clear APOE, we wanted to study how the neurons recover from APOE exposure. To understand this, we added a 20-minute recovery period following 20-minute APOE4 conditioned media exposure. When we tested the neurons after 20 minutes of recovery, phosphorylation of eEF2 was significantly reduced (**Fig S5D**), implying that the APOE4 mediated effect could be recovered. To further study the rate of recovery and their differences between the APOE isoforms, we treated the neurons with either APOE3 or APOE4 conditioned media for 1 minute (a time point where both APOE3 and APOE4 induced a robust increase in eEF2 phosphorylation - **Fig 5C**), followed by a recovery of 5, 10 and 20 minutes (**Fig 5F**). Interestingly, after 1-minute treatment followed by 5 minutes of recovery, the phosphorylation of eEF2 was still significantly higher in neurons exposed to APOE4 compared to APOE3 treated neurons (**Fig 5F**). But, by 10 minutes and 20 minutes of recovery, both APOE3 and APOE4 treated neurons had reached the same level of eEF2 phosphorylation. This indicated that the rate of recovery from translation inhibition was significantly slower in APOE4 treated neurons. Finally, to test the long-term effect of APOE, we treated the neurons with APOE conditioned media for 24 hours and measured the phosphorylation status of eEF2 (**Fig S5B**). At 24 hours, phosphorylation of eEF2 was significantly higher in neurons treated with APOE4 conditioned media compared to APOE3 or APOE KO conditioned media (**Fig S5B**). Similarly, phosphorylation of ERK was also significantly higher in neurons treated with APOE4 conditioned media for 24 hours (**Fig S5C**). This indicated that the chronic exposure of APOE4 also caused translation inhibition. In summary, whilst both APOE3 and APOE4 induced a robust translation inhibition at 1 minute similar to NMDAR stimulation, their responses followed different temporal trajectories. The recovery from translation inhibition was slower in APOE4 treated neurons indicating the involvement of additional components in the APOE4 induced response.

### The translation response of APOE and NMDARs are closely linked to their calcium signature

An important feature of APOE4 induced translation inhibition was that it resembled NMDAR stimulation but also abolished the actual NMDAR response. Since the NMDAR translation response is primarily due to the influx of calcium into the neurons^44^, we tested the effect of extracellular calcium in the APOE4 induced translation response. We treated the neurons with recombinant APOE3 or APOE4 protein for 20 minutes in ACSF medium with or without calcium (**Fig 6A)**. As shown in the previous figures, incubation with APOE4 recombinant protein for 20 minutes resulted in a significant increase in eEF2 phosphorylation in comparison to APOE3 (**Fig 6A**), but the absence of calcium in the media significantly reduced the APOE4 induced increase in the phosphorylation of eEF2. Though the absence of calcium showed a trend of decrease in eEF2 phosphorylation in the presence of APOE3 as well, it was not significant (**Fig 6A**). The absence of calcium in the control media or ACSF did not affect the phosphorylation of eEF2 (**Fig S6A**). Thus, calcium influx had an important role in APOE4 mediated translation inhibition. Hence, we wanted to understand the calcium profiles of APOE3 and APOE4, and also compare it to the calcium profile of NMDAR stimulation. To study this, we measured the changes in cytosolic calcium in neurons on NMDA or APOE addition for 5 minutes using Fluo4-AM dye.

**Figure 6.**
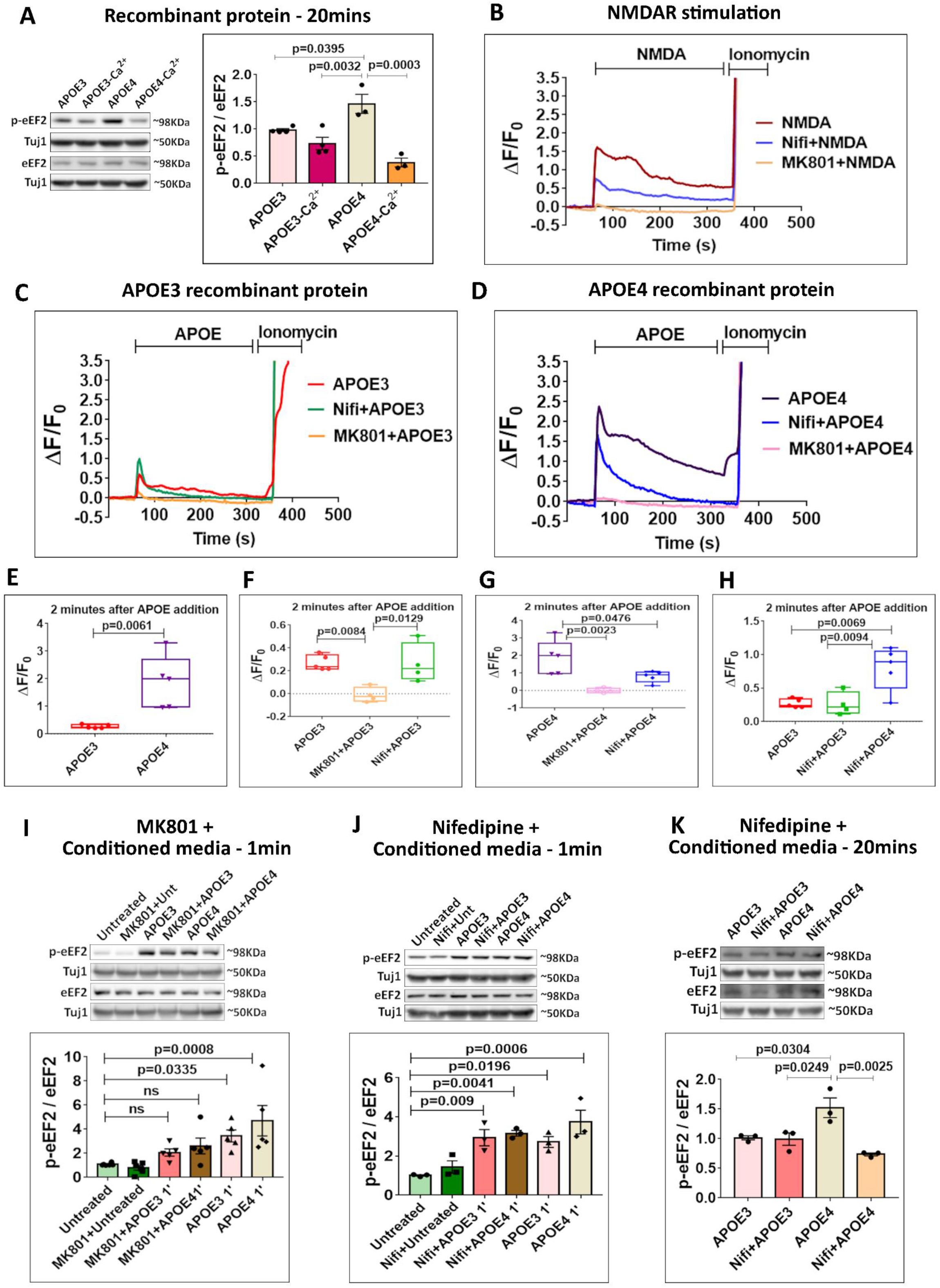
The APOE mediated translation response is regulated by calcium signature and its sources. **A** - Rat primary cortical neurons (DIV15) were treated with recombinant APOE3/APOE4 protein (15nM) for 20minutes in the presence or absence of extracellular calcium (ACSF with or without calcium) and probed for the phosphorylation of eEF2. Left - representative immunoblots indicating levels of phospho-eEF2, eEF2 and Tuj1; Right - graph indicating ratio of phospho-eEF2 to eEF2 normalized to Tuj1. Data is represented as mean +/- SEM. N=3-4, One-way ANOVA (p=0.0004) followed by Tukey’s multiple comparison test. Rat primary cortical neurons (DIV15) were subjected to calcium imaging for 5 minutes. The graphs represent the time trace for the change in Flou4-AM fluorescence compared to initial fluorescence (ΔF/F0) under the following conditions - **B-** NMDAR stimulation (20μM), Nifedipine (50μM) pre-treatment followed by NMDAR stimulation (20μM) and MK801 (25μM) pre-treatment followed by NMDAR stimulation (20μM). **C** - APOE3 treatment (15nM), Nifedipine (50μM) pre-treatment followed by APOE3 addition (15nM) and MK801 (25μM) pre-treatment followed by APOE3 addition (15nM). **D** - APOE4 treatment (15nM), Nifedipine (50μM) pre-treatment followed by APOE4 addition (15nM) and MK801 (25μM) pre-treatment followed by APOE4 addition (15nM). **E, F, G, H** – Box plots represent the quantification of the change in Flou4-AM fluorescence (ΔF/F0) after 2minutes of APOE addition. N=4-5 experiments, each experiment has the average value of 40-50 neurons. E - Unpaired Student’s t-test; F - One-way ANOVA (p=0.0058) followed by Tukey’s multiple comparison test; G - One-way ANOVA (p=0.0029) followed by Tukey’s multiple comparison test; H - One-way ANOVA (p=0.0038) followed by Tukey’s multiple comparison test. **I** - Rat primary cortical neurons (DIV15) were treated with MK801 (25μM) along with APOE3 or APOE4 (10-15nM) conditioned media for 1 minute and probed for phosphorylation of eEF2. Top - representative immunoblots indicating levels of phospho-eEF2, eEF2 and Tuj1; Bottom - graph indicating ratio of phospho-eEF2 to eEF2 normalized to Tuj1. Data is represented as mean +/- SEM. N=5. One-way ANOVA (p=0.0006) followed by Dunnett’s multiple comparison test. **J** - Rat primary cortical neurons (DIV15) were treated with Nifedipine (50μM) along with APOE3 or APOE4 (10-15nM) conditioned media for 1 minute and probed for phosphorylation of eEF2. Top - representative immunoblots indicating levels of phospho-eEF2, eEF2 and Tuj1; Bottom - graph indicating ratio of phospho-eEF2 to eEF2 normalized to Tuj1. Data is represented as mean +/- SEM. N=3. One-way ANOVA (p=0.001) followed by Dunnett’s multiple comparison test. **K** - Rat primary cortical neurons (DIV15) were treated with Nifedipine (50μM) along with APOE3 or APOE4 (10-15nM) conditioned media for 20 minutes and probed for phosphorylation of eEF2. Top - representative immunoblots indicating levels of phospho-eEF2, eEF2 and Tuj1; Bottom - graph indicating ratio of phospho-eEF2 to eEF2 normalized to Tuj1. Data is represented as mean +/- SEM. N=3. One-way ANOVA (p=0.0036) followed by Tukey’s multiple comparison test.

On NMDAR stimulation, there was a rapid influx of calcium and the increase in cytosolic calcium levels sustained up to 5 minutes (**Fig 6B**). NMDAR antagonist MK801 completely abolished NMDAR mediated calcium influx (**Fig 6B**), confirming the specificity of the response. It was previously reported that L type-voltage gated calcium channels (L-VGCCs) are essential for the sustenance of NMDAR mediated calcium^52^. Inhibition of L-VGCCs using Nifedipine did not block the influx of calcium on NMDAR stimulation, but it eliminated the later part of the sustained calcium response (**Fig 6B**). This verified that L-VGCCs is an important contributor to NMDAR induced calcium influx. The addition of recombinant APOE3 protein led to a small burst of calcium (**Fig 6C**) whereas the addition of recombinant APOE4 protein caused a robust and sustained increase in the cytosolic calcium which was substantially bigger than APOE3 (**Fig 6C and 6D**). The change in fluorescent intensity calculated after APOE addition also showed that APOE4 caused a significant increase in calcium levels compared to APOE3, both at the 1-minute and 2-minute time point (**Fig 6E and S6B**). MK801 treatment abolished the calcium influx for both APOE3 and APOE4 addition (**Fig 6C and 6D**) indicating that NMDARs were the primary source of calcium. Interestingly, treatment with L-VGCC specific inhibitor Nifedipine prevented the APOE4 mediated sustenance of calcium while it did not affect the APOE3 mediated calcium response (**Fig 6C and 6D**). This indicated that APOE4 exposure caused a sustained increase in cytosolic calcium levels which was initiated by calcium through NMDARs but the latter part of sustenance was primarily contributed by L-VGCCs.

Further quantification of the change in fluorescent intensities showed that APOE3 exposure led to a small increase in the cytosolic calcium which was rescued by MK801 both at 1-minute and 2-minute time points, but Nifedipine was not able to rescue this at either of the time points (**Fig 6F and S6C**). This showed that the APOE3 calcium response was mediated by NMDARs only. In the case of APOE4, while Nifedipine pre-treatment reduced the calcium levels on APOE4 addition at 1-minute, it was still significantly higher compared to MK801 pre-treatment (**Fig S6D**). But, by the 2-minute time point, Nifedipine also significantly reduced the APOE4 mediated calcium levels (**Fig 6G**). This indicated that the contribution of L-VGCCs was more for APOE4 mediated calcium response, particularly from 2-minute onwards, rather than the earlier 1-minute timepoint. Finally, even in the presence of Nifedipine, the increase in cytosolic calcium levels caused by APOE4 was significantly higher than APOE3 both at 1-minute and 2-minutes, indicating that the initial NMDAR component was also higher in the case of APOE4 (**Fig 6H and S6E**). In summary, both APOE3 and APOE4 caused initial calcium influx through NMDARs, but the extent of NMDAR mediated calcium entry was higher in the case of APOE4. At a later phase, APOE4 also caused an influx of calcium through L-VGCCs which did not happen in the case of APOE3. We also observed that chronic exposure (24 hours) of APOE4 in the form of recombinant protein (**Fig S6F**) or conditioned media (**Fig S6G**) led to a significant increase in the cytosolic calcium levels in neurons. This sustained calcium increase was likely to be the cause of aberrant translation inhibition and the loss of synaptic activity mediated translation response in APOE4 treated neurons.

Further, we established the link between the different sources of calcium and the translation response generated by APOE. We tested the effect of 1-minute treatment of APOE3 and APOE4 conditioned media on eEF2 phosphorylation in the presence of MK801 (open channel NMDAR inhibitor). As shown previously (**Fig 5**), 1-minute treatment of APOE3 or APOE4 conditioned media led to a significant increase in eEF2 phosphorylation (**Fig 6I**). However, in the presence of MK801, the phosphorylation of eEF2 on 1-minute exposure to APOE3 and APOE4 was not significantly different from the untreated neurons (**Fig 6I**). This indicated that at 1 minute both APOE3 and APOE4 caused translation inhibition due to the influx of calcium through NMDARs. Since we observed that the later part of APOE4 mediated calcium was through L-VGCCs, we tested whether blocking L-VGCCs would rescue the translation defect caused by APOE4. Preincubation of neurons with Nifedipine did not have any effect on the APOE3 or APOE4 induced translation response at 1-minute as measured by the phosphorylation of eEF2 (**Fig 6J**). This corroborated the finding that L-VGCCs did not contribute to the APOE3 calcium response, and L-VGCCs did not have a role in APOE4 calcium response at 1-minute. However, at 20 minutes, Nifedipine significantly reduced APOE4 mediated increase in eEF2 phosphorylation (**Fig 6K**) confirming the role of L-VGCCs in the later part of the APOE4 mediated defect. MK801 or Nifedipine alone did not have any effect on eEF2 phosphorylation in neurons exposed to APOE KO conditioned media (**Fig S6H and S6I**). Hence, the translation inhibition caused by APOE4 has an early phase contributed by calcium through NMDARs while the sustenance of this response was due to the calcium through L-VGCCs. The perturbation of calcium homeostasis disrupted the basal and NMDAR mediated translation response in APOE4 treated neurons.

## Discussion

In this study, we showed that APOE4 inhibits protein synthesis in neurons and isolated synaptic compartments. We tested this by using different sources of APOE like APOE from stem cell conditioned media, astrocyte conditioned media, and recombinant protein. It is also important to note that we have used APOE at physiologically relevant low nanomolar concentrations (10nM)^53,54^. An important aspect of our study was that we were able to identify the distinct temporal profiles of protein synthesis responses generated by APOE3 and APOE4. This gave us new insights into how APOE3 and APOE4 lead to translation inhibition at an earlier time point (1-minute) and the key difference existed in their rate of recovery from the inhibition, where APOE4 mediated recovery was slower. The extent of activation of eEF2 phosphorylation was one of the major differences between APOE3 and APOE4. Additionally, we showed that the temporal differences in the responses generated by the APOE isoforms also contribute significantly to making APOE4 harmful. This brought in an interesting perspective towards the differences between the APOE isoforms. Our results indicating the long-lasting effect of APOE4 on protein synthesis inhibition is of specific importance when placed in the context of synaptic defects caused by APOE4. Similar to the studies which show protein synthesis as an early defect in familial AD models^15,16,19^, our study raises the possibility of translation dysregulation as an early defect in APOE4 mediated pathology as well.

In neurons, protein synthesis is regulated in a complex and dynamic manner, and heavily modulated downstream of synaptic activity^3,4^. This tight spatio-temporal regulation of synaptic translation is well studied for NMDAR stimulation^44,49,55,56^. Initially, NMDAR stimulation leads to a global protein synthesis inhibition which changes to a phase of translation activation with time^44,49,50^. The interesting finding was that both APOE3 and APOE4 treatment appeared to mimic the translation inhibition phase of NMDAR stimulation. While APOE3 recovered from the inhibitory phase to basal state faster than APOE4, neither of the APOE isoforms activated translation at a later time point (20-minute) like NMDAR stimulation. Thus, the primary difference lied in the distinct temporal profiles of APOE treatment and NMDAR stimulation. Moreover, these results call attention to the specificity of the translation response downstream of synaptic stimulation.

We have previously shown that the inhibitory phase of NMDAR translation response involved the translation activation of a specific subset of mRNAs like PTEN and PSD95^49^. APOE4 treatment (but not APOE3 treatment), not only mimicked the translation inhibition phase of NMDAR stimulation, but it also caused the translation activation of PTEN and PSD95 mRNAs. Since we confirmed that APOE4 exposure mimicked the specific translation response of NMDAR stimulation, we wanted to test the response of APOE treated neurons to physiological NMDAR stimulation. In the presence of APOE4, the NMDAR specific protein synthesis response was lost, both in terms of translation inhibition and candidate-specific translation activation. However, APOE3 treatment did not affect the NMDAR protein synthesis response. Another interesting finding was that NMDAR stimulation in the presence of APOE4 led to a stress response phenotype by causing an inhibition of translation initiation and eIF2α phosphorylation. This indicated that along with the basal translation, activity mediated protein synthesis response which is salient in shaping synaptic plasticity was also impaired in APOE4 treated neurons. This further implicated that disruption of synaptic signaling was an important player in APOE4 pathology.

The specificity of the signal generated downstream of NMDARs is linked to their feature of high calcium permeability^57–59^. Stimulation with NMDARs leads to a robust increase in intracellular calcium contributed by different sources – NMDA receptors, L type-Voltage Gated Calcium Channels (L-VGCCs), and internal calcium stores^52,57,60–63^. Many studies have outlined the idea that the activation of these different sources of calcium is spatially and sequentially regulated^57,60,62,64^. NMDARs lead to the first burst of calcium which further activates L-VGCCs, followed by internal stores through calcium-induced calcium release (CICR)^52,60–62^. Thus, synaptic stimulation generates a distinct calcium signature which is also tightly regulated spatially and temporally. Calcium being one of the most important secondary messengers, many kinases and phosphatase are sensitive to calcium^57,65^, directly influencing signaling pathways that regulate protein synthesis^57^. Previous studies have outlined the contribution of calcium in causing the inhibitory phase of NMDAR translation response^44,55^. Thus, we hypothesize that the distinct translation response generated downstream of synaptic stimulation could be a result of the specific calcium signature generated by them. Hence, we explored the link between protein synthesis response and calcium homeostasis in the presence of APOE isoforms as a possible explanation for APOE4 mediated synaptic translation defects.

APOE was previously reported to cause an influx of calcium in neurons and this has been linked to the activation of NMDARs^39,48,66–68^. Accordingly, we observed that the APOE4 mediated translation inhibition (phosphorylation of eEF2) was indeed dependent on calcium. Thus, we assessed the calcium signatures generated by APOE3 and APOE4 and identified their sources of calcium influx. APOE3 led to a small burst of calcium where the source of calcium was NMDARs. However, APOE4 caused a sustained increase in intracellular calcium where the contributing sources were both NMDARs and L-VGCCs. This showed that both APOE3 and APOE4 activated NMDARs and caused calcium influx through it. The NMDAR antagonist MK801 completely blocked the calcium influx caused by both APOE isoforms, further highlighting that NMDARs were the primary source of calcium entry. Consecutively, MK801 was also able to prevent the APOE3 and APOE4 mediated translation inhibition at the early time point (1-minute). APOE receptors and NMDARs are known to interact with each other^33,35,39^. Activation of the ERK signaling pathway by APOE is also shown to be dependent on NMDAR activation^39,69,70^. Additionally, the Dab1-SFK signaling downstream of APOE receptors is shown to phosphorylate and activate NMDARs^39,71^. Though several studies have strengthened the influence of APOE receptor-associated signaling on NMDARs^72^, the exact mechanism behind APOE mediated calcium influx through NMDARs is yet to be understood. Another possible mechanism of APOE mediated NMDAR activation could be through the production of Aβ. APOE4 has been shown to cause an increase in Aβ production^37,73^ and Aβ is shown to directly activate NMDARs^74^. Apart from the involvement of NMDARs, our results highlight the importance of L-VGCCs in sustaining the calcium increase on APOE4 treatment. L-VGCC antagonist Nifedipine did not block the initial calcium entry on APOE4 treatment but its sustenance, further supporting the idea of sequential calcium entry where NMDARs were the first source and L-VGCCs were the latter. However, it is important to note that the initial NMDAR calcium component was also higher in the case of APOE4 compared to APOE3. The higher calcium influx through NMDARs could be the plausible explanation for the L-VGCC activation by APOE4 only. Corresponding to the calcium profile, Nifedipine could not prevent the APOE4 mediated translation inhibition at the earlier time point (1-minute), while it successfully rescued it at the later time point (20-minute). These results support the idea of distinct calcium signatures generated by the sequential activation of different sources. It also emphasizes that the different calcium sources and signatures clearly lead to distinct translation responses. Hence, it highlights the importance of studying APOE4 mediated calcium dysregulation with a focus on spatio-temporal calcium signatures rather than individual sources.

As stated previously, one of the key features of the NMDAR mediated translation profile was the later phase of global protein synthesis activation which did not occur with APOE3 and APOE4. We are yet to understand the role of calcium-mediated regulation in this phase of global translation activation. However, we hypothesized that the internal calcium release (CICR) could have an important contribution to this aspect. The stimulation of NMDARs leading to L-VGCC activation is reported to initiate CICR from the ER stores^52^. The activation of CICR is shown to have a feedback inhibition on the calcium entry through L-VGCCs and NMDARs^52^. Hence, the intracellular calcium increase downstream of NMDAR activation is contributed by calcium from external sources (NMDARs and L-VGCCs) followed by calcium from internal sources (ER stores), likely to occur in distinct phases which are sequential and spatially separated. We hypothesized that these distinct calcium phases/sources could be coupled to the distinct translation phases observed on NMDAR activation. Along the lines of this hypothesis, we propose that the release of calcium from internal sources could be the key difference between the APOE isoforms and NMDAR stimulation, explaining the absence of the global translation activation phase on APOE treatment.

It is important to note that we focused on the effect of APOE on translation at the post-synapse as protein synthesis regulation is better understood at the post-synapse. Our model system was the cultured primary cortical neurons. Hence it lacked the APOE clearance mechanisms which exist physiologically. Thus, we speculate that many of the phenotypes we observed might be exaggerated and it would be much more subtle in the brain. Previous studies have reported the effects of APOE4 treatment on transcription, especially that of PSD95 mRNA^38^. We did not observe any change in the PSD95 steady-state mRNA levels with our system of treatment and APOE concentrations. Since we have used the phosphorylation status of eEF2 as a readout for most of our assays, we checked the mRNA levels of eEF2 kinase (eEF2K) and eEF2 phosphatase (PP2a). These were not affected by APOE treatment in our experimental conditions (data not shown). But, the effect of APOE on the protein levels / the activity of the kinases and phosphatases are unknown, and could contribute to the changes observed in eEF2 phosphorylation. However, we have used many direct readouts like FUNCAT and polysome profiling to validate the results on protein synthesis regulation.

With all the above points in consideration, we propose the model (**Figure 7**) where we explain the specific calcium and translation signatures downstream of NMDAR stimulation and APOE treatment. NMDAR stimulation leads to calcium influx through NMDARs and L-VGCCs leading to its distinct translation response of early phase inhibition (corresponding to increased eEF2 phosphorylation) followed by a later phase activation (corresponding to decreased eEF2 phosphorylation) (**Figure 7A**). APOE3 causes a short burst of calcium influx through NMDARs alone causing an early phase translation inhibition that recovers to the basal level. Hence, APOE3 treatment followed by NMDAR stimulation does not affect the NMDA mediated translation response (**Figure 7B**). APOE4 causes a robust and sustained influx of calcium through NMDARs and L-VGCCs. This leads to a sustained translation inhibition phase which recovers much slower than APOE3. Hence, in the background of APOE4 treatment, NMDAR stimulation leads to a stress response causing further reduction of protein synthesis (**Figure 7C**).

**Figure 7.**
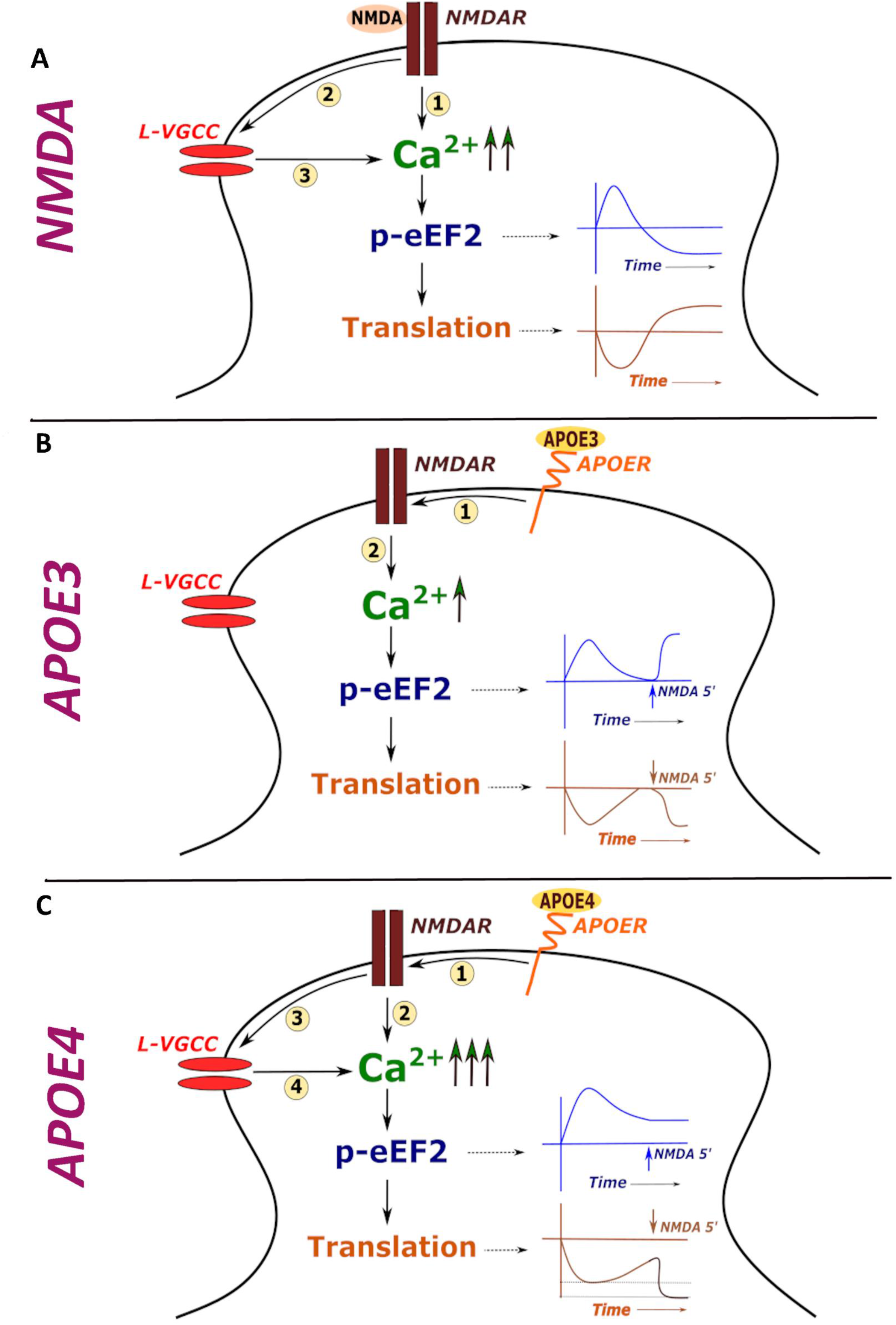
Model illustrating calcium signature and corresponding protein synthesis regulation kinetics downstream of NMDAR stimulation and APOE treatment. A - Stimulation of NMDA receptors leads to influx of calcium through NMDARs and L-VGCCs which generates a specific temporal profile of eEF2 phosphorylation. This leads to an initial inhibition of global protein synthesis followed by translation activation. B - Exposure to APOE3 activates NMDARs leading to a short burst of calcium through it. This leads to an acute increase in eEF2 phosphorylation which eventually recovers to basal levels. Global translation also follows a similar temporal profile of initial decrease followed by recovery. Hence, the NMDA activity mediated translation response is unaffected in APOE3 treated neurons. C - Exposure to APOE4 activates both NMDARs and L-VGCCs leading to a huge and sustained increase in calcium levels. This leads to the sustained increase in eEF2 phosphorylation as well as global translation inhibition. Hence, the NMDA activity mediated response is perturbed, potentially causing a stress-response phenotype in APOE4 treated neurons.

Thus, our work identifies the translation defect caused by APOE4 and links it to the calcium signature generated by it. Additionally, it brings forth a new perspective towards protein synthesis regulation brought about by the distinct calcium signatures and their sources. APOE4 is shown to cause synaptic dysfunction which leads to deficits in learning and memory. Now, many studies are indicating how APOE4 could cause AD phenotype independent of classical familial mutations of AD^37,73^. We propose that protein synthesis dysregulation could be a central component responsible for synaptic and cognitive defects in the APOE4 mediated pathology.

## Materials and Methods

### Ethics Statement

All animal work was carried out in accordance with the procedures approved by the Institutional Animal Ethics Committee (IAEC) and the Institutional Biosafety Committee (IBSC), InStem, Bangalore, India. All rodent work was done with Sprague Dawley (SD) rats. Rats were kept in 20–22 temperature, 50–60 relative humidity, 0.3 μm HEPA-filtered air supplied at 15–20 ACPH, and 14-h/10-h light/dark cycle maintained.

All the human stem cell work was carried out in accordance with and approval from the Institutional Human Ethics Committee, Institutional Stem Cell Committee, and Institutional Biosafety Committee at inStem, Bangalore, India.

### Rat primary neuronal cultures

Primary neuronal cultures were prepared from cerebral cortices of Sprague-Dawley rat embryos (E18.5) as previously published by our lab^44,49,75^. Briefly, the cortex tissue was trypsinized and homogenized using 0.25% trypsin. The dissociated cells were plated on pre-coated Poly-L-Lysine (P2636, Sigma) dishes at a density of 40000-50000 cells/ cm^2^ for biochemistry experiments and 30000-40000 cells/cm^2^ for imaging-based experiments. The plates were coated with Poly-L-Lysine solution (0.2mg/ml) made in borate buffer (pH 8.5) for 5-6 hours and excess solution was washed with water. For imaging experiments, the neurons were plated on nitric acid-treated coverslips which were coated with Poly-L-Lysine. For all the experiments, the neurons were initially plated on Minimum Essential Media (MEM, 10095080, ThermoFisher Scientific) supplemented with 10% FBS to aid their attachment. After 3 hours in MEM, the media was changed to Neurobasal (21103049, ThermoFisher Scientific) supplemented with B27 (17504044, ThermoFisher Scientific) and 1X Glutamax. The neurons were maintained in culture for 15-20 days at 37°C, 5% CO_2_ conditions by supplementing Neurobasal media every 5-6 days.

### iPSC maintenance and conditioned media collection

The iPSCs (APOE KO, APOE 3/3, and APOE 4/4) were obtained from Bioneer A/S, Denmark. Briefly, the iPSCs from an 18-year old male of APOE 3/4 genotype were subjected to CRISPR-Cas9 gene editing to obtain isogenic iPSC lines of APOE KO, APOE 3/3, and APOE 4/4 genotypes^42^. The iPSCs were maintained in mTeSR1 complete media (72232, Stem Cell Technologies) at 37°C, 5% CO_2_ conditions. A mixture of 1mg/ml Collagenase IV (17104019, ThermoFisher Scientific), 0.25% Trypsin, 20% Knock-Out Serum (10828028, ThermoFisher Scientific), 1mM Calcium Chloride (made in PBS) was used to dissociate the iPSCs for passaging.

Once the iPSCs reached about 50% confluency, the media was changed from mTeSR1 to Neurobasal + Glutamax. The iPSCs were maintained in Neurobasal based media for 48 hours. After 48 hours, the conditioned media was given a short spin at 1000 rcf for 2 minutes to remove the cell debris and subjected to ELISA (ab108813, Abcam) to estimate the amount of APOE secreted by iPSCs.

### Synaptoneurosome preparation

Synaptoneurosomes were prepared from cortices of P30 Sprague-Dawley rats as previously published^49,76,77^. Briefly, cortices were homogenized at 4°C in 10 volumes of the synaptoneurosome buffer (118mM NaCl, 5mM KCl, 1.2mM MgSO_4_, 2.5mM CaCl_2_, 1.53mM KH_2_PO_4_, 212.7mM Glucose, 1X Protease Inhibitor Cocktail, pH 7.5). The homogenate was filtered through three 100μM nylon filters (NY1H02500, Merck Millipore) and one 11μM nylon filter (NY1102500, Merck Millipore). The filtrate was centrifuged at 1500 rcf for 15 minutes at 4°C. The pellet obtained was resuspended in 1.2ml synaptoneurosome buffer and used for APOE treatment.

### Treatment with APOE and other drugs

a. **Treatment of primary neurons** - The primary neurons were treated with APOE from conditioned media or recombinant APOE protein (350-02, 350-04, Peprotech). For conditioned media treatment, the spent neuronal media was mixed with the APOE conditioned media in the ratio 1: 1 such that the final APOE concentration used to treat the neurons was 10-20nM. In case of recovery, the conditioned media mixture was removed and the spent neuronal media was added for recovery. For recombinant protein treatment, the neurons were treated with 15nM of recombinant APOE3 or APOE4 protein in neurobasal media. In the experiments with no calcium in the external media, the neurons were treated with 15nM recombinant APOE3 or APOE4 protein in Artificial Cerebrospinal Fluid (ACSF – 120mM NaCl, 3mM KCl, 1mM MgCl_2_, 3mM NaHCO_3_, 1.25mM NaH_2_PO_4_, 15mM HEPES, 30mM glucose, pH 7.4) with or without calcium (2mM CaCl_2_). For stimulation of NMDAR receptors, 20μM NMDA (0114, Tocris) was added during the last 5 minutes of the 20 minutes treatment. For the pre-treatment with other drugs, Nifedipine (50μM) (N7634, Sigma), MK801 (25μM) (0924, Tocris), or RAP (200nM) (553506-M, Sigma) were treated for 10 minutes before the addition of conditioned media mixture. After the treatment, the cells were lysed in buffer containing 20mM Tris-HCl, 100mM KCl, 5mM MgCl_2_, 1% Nonidet P-40 (NP40), 1mM dTT, 1X Protease Inhibitor Cocktail, RNase inhibitor, 1X Phosphatase Inhibitor and centrifuged at 20000g, 4°C for 20 minutes. The supernatant was either denatured in SDS-dye for Western blotting or in Trizol LS for RNA isolation.
b. **Treatment of synaptoneurosomes** - The resuspended synaptoneurosomes were treated with 20nM recombinant APOE3 or APOE4 protein for 20 minutes at 37°C with mixing at 350rpm. For stimulation of NMDA receptors in the syanptoneurosomes, NMDA (40μM) was added during the last 5 minutes of the 20 minutes treatment. After the treatment, the synaptoneurosomes were given a short spin, the pellet was resuspended in lysis buffer (20mM Tris-HCl, 100mM KCl, 5mM MgCl_2_, 1% Nonidet P-40 (NP40), 1mM dTT, 1X Protease Inhibitor Cocktail, RNase inhibitor, 1X Phosphatase Inhibitor) and centrifuged at 20000g, 4°C for 20 minutes. The supernatant was denatured in SDS-dye and subjected to Western blotting.

### Mouse primary neuronal culture

Primary mouse neurons were generated from cortical and hippocampal tissue of wild-type embryos (E15-E17). Wild-type mouse embryos were obtained from pregnant B6.Cg-Tg (APPswe,PSEN1dE9)85Dbo/Mmjax mice (APP/PS1) mice, a widely used transgenic model of AD-like amyloidosis. The protocol used to generate primary neurons was previously described^78^. In short, cortices and hippocampi were dissected, incubated in 0.25% trypsin for 15 minutes, and dissociated in Dulbecco’s modified Eagle medium (DMEM) containing 10% Fetal Bovine Serum (FBS) (Gibco, 1008214) and 1% Penicillin/Streptomycin (ThermoFisher Scientific, SV3001). Dissociated primary cells were seeded on poly-D-lysine coated plates and maintained until 15 days in vitro in Neurobasal medium with B27 supplement, 1.4 mM L-glutamine, and 1% Penicillin/Streptomycin.

### Mouse primary astrocyte culture

Primary astrocyte cultures were obtained from ApoE3 KI (B6.Cg-Apoe^em2(APOE*)Adiuj/J^, the Jackson Laboratory, #029018), ApoE4 KI (B6(SJL)-Apoe^tm1.1(APOE*4)Adiuj/J^, the Jackson Laboratory, #027894) and ApoE KO (B6.129P2-Apoe^<tm1Unc>/J^, the Jackson Laboratory, #002052)^79^ mouse brains. Cortical and hippocampal tissue was dissected from post-natal pups (P1-P3). After dissection, the brain tissue was incubated in 0.25% trypsin for 15 minutes. The tissue was dissociated into cells in DMEM containing 5% FBS (Gibco, 1008214) and 1% Penicillin/Streptomycin (ThermoFisher Scientific, SV3001) using soft plastic Pasteur pipettes. Primary cells were seeded at a concentration of 500,000 cells on poly-D-lysine coated T75 flasks. 3-5 hours after seeding and culturing in DMEM medium containing 5% FBS and 1% P/S, the medium was replaced by AstoMACS medium (Miltenyi Biotec, 130-117-03) with 0.5 mM L-glutamine. AstroMACS medium of the primary astrocytes was replaced every 2-3 days with fresh AstoMACS medium. When at least 80% confluence of the primary astrocyte cultures was reached (around DIV12), the astrocytes were cultured in Neurobasal medium supplemented with B27 supplement, 1% Penicillin/Streptomycin, and 1.4 mM L-glutamine for 48 hours for the collection of astrocyte conditioned medium. The conditioned astrocyte medium was used for the APOE treatment of the wild-type mouse primary neurons. The primary mouse neuron and astrocyte experiments were approved by the Ethical Committee for animal research at Lund University.

### FUNCAT (Fluorescent non-canonical amino acid tagging)

For metabolic labeling, DIV15 neurons were incubated in Methionine-free DMEM for 45 minutes. Followed by this, the neurons were treated with L-azidohomoalanine (AHA, 1μM) (1066100, Click Chemistry tools) for 30 minutes in Met-free DMEM (21013024, ThermoFisher Scientific). Followed by this, they were treated with 15nM APOE3 or APOE4 recombinant protein in the same media (Met-free DMEM with AHA). For stimulation of NMDAR receptors, NMDA (20μM) was added during the last 5 minutes of the 20 minutes treatment. After the treatment, the coverslips were given one wash with 1X PBS and fixed with 4% PFA. After 15 minutes of fixing, they were washed thrice with 1X PBS. The neurons were permeabilized for 10 minutes with 0.3% Triton X-100 solution prepared in TBS_50_ (50mM Tris, 150mM NaCl, pH 7.6). The permeabilized neurons were subjected to blocking for 1 hour with a mixture of 2% Bovine Serum Albumin (BSA) and 2% Fetal Bovine Serum (FBS) prepared in TBS_50_t (TBS50 with 0.1% Triton X-100). After blocking, the neurons were subjected to the FUNCAT reaction for 2 hours where the newly-synthesized AHA incorporated proteins were tagged with an alkyne-fluorophore Alexa-Fluor 555 through click reaction (C10269, CLICK-iT cell reaction buffer kit, Click Chemistry Tools). After 3 washes with TBS_50_t, the neurons were stained with MAP2 antibody (1:1000 dilution prepared in blocking buffer incubated overnight at 4°C) for detection of neuronal cells. This was followed by secondary antibody staining with Alexa-Fluor 488 (1:500 dilution prepared in blocking buffer, incubated for 1 hour at room temperature) (A-11008, ThermoFisher Scientific) to visualize MAP2. The coverslips were mounted with Mowiol and imaged on Olympus FV300 confocal laser scanning inverted microscope with 60X objective. The pinhole was kept at 1 Airy Unit and the optical zoom at 2X to satisfy Nyquist’s sampling criteria in XY direction. The objective was moved in Z-direction with a step size of 1μM (~8-9 Z-slices) to collect light from the planes above and below the focal plane. The image analysis was performed using FIJI software and the maximum intensity projection of the slices was used for quantification of the mean fluorescent intensities. The mean fluorescent intensity of FUNCAT channel was normalized to the MAP2 channel for comparison between different APOE treatment conditions.

### Immunostaining

The APOE iPSCs grown on 4-well dishes were subjected to immunostaining of the pluripotency markers OCT4 and NANOG. The iPSCs were fixed with 4% PFA which was followed by 3 washes with 1X PBS. This was followed by permeabilization with 0.3% Triton X-100 made in TBS50. This was followed by 1 hour blocking with 2% BSA and 2% FBS prepared in TBS50T (with 0.1% Triton X-100). They were incubated with the primary antibody (prepared in blocking buffer) overnight at 4°C. This was followed by 3 washes with TSB50T and 1-hour incubation with the secondary antibody (prepared in blocking buffer) at room temperature. After 3 washes with TBS50T, the iPSCs were subjected to post-fixing with 4% PFA. This was followed by 3 washes with 1X PBS and mounted with Mowiol. The 4 well plates were imaged on Olympus IX73 inverted fluorescence microscope with 20X objective.

### Western blotting

The APOE treated neuron lysates or synaptoneurosome lysates were subjected to western blotting analysis. Briefly, the denatured lysates were run on 10% resolving and 5% stacking acrylamide gels and subjected to overnight transfer onto PVDF membrane. The blots were subjected to blocking for 1 hour at room temperature using 5% BSA prepared in TBST (TBS with 0.1% Tween-20). This was followed by primary antibody (prepared in blocking buffer) incubation for 2-3 hours at room temperature. HRP (Horseradish peroxidase) tagged secondary antibodies were used for primary antibody detection. The secondary antibodies (prepared in blocking buffer) were incubated with the blots for 1 hour at room temperature. Three washes of 5-10 minutes each were given after primary and secondary antibody incubation using TBST solution. The blots were subjected to chemiluminescent based detection of the HRP tagged proteins. For the analysis of eEF2 phosphorylation and ERK phosphorylation, the samples were run in duplicates where one set was used to probe for the phospho-proteins (p-eEF2 and p-ERK), and the other set was used to probe for the total proteins (eEF2 and ERK). In each set, the loading control used was Tuj1. In every set, eEF2 and ERK were probed on the same blot. Hence, the Tuj1 used to normalize eEF2 and ERK was the same in a given set. Similarly, p-eEF2 and p-ERK were probed on the same blot for a given set. Hence, Tuj1 used to normalize them was also the same for the given set. For eIF2 phosphorylation analysis, the samples were run in duplicates (one set for phospho-protein, one set for total protein). Tuj1 was used as the normalizing control for each blot. For PTEN and PSD95 level analysis, PTEN and PSD95 were probed on the same blot and Tuj1 was used as the normalizing control for each set. For RPLP0 and RPS6 distribution analysis, the individual polysome fractions were run and used for percentage distribution quantification. The details of the antibodies used and their dilutions are given in the table at the end. All the western blot quantifications were performed using densitometric analysis on ImageJ software.

### Polysome profiling

The DIV15 neurons (~1.5 to 2 million cells) were treated with APOE3 or APOE4 iPSC conditioned media for 20 minutes. For stimulation of NMDARs, NMDA (20μM) was added during the last 5 minutes of the 20 minutes treatment. After the treatment, the cells were lysed in buffer containing 20mM Tris-HCl, 100mM KCl, 5mM MgCl_2_, 1% Nonidet P-40 (NP40), 1mM dTT, 1X Protease Inhibitor Cocktail, RNase inhibitor, 0.1mg/ml Cycloheximide (C7698, Sigma), 1X Phosphatase Inhibitor and centrifuged at 20000g, 4°C for 20 minutes. The supernatant was loaded on 15%-45% linear sucrose gradient prepared in buffer containing 20mM Tris-HCl, 100mM KCl, 5mM MgCl_2_, 0.1mg/ml Cycloheximide. The gradients loaded with the lysates were subjected to ultracentrifugation at 39000rpm, 4°C for 1.5 hours. 1ml fractions were collected from the gradient after the spin (11 fractions in total); as the fractions were collected, they were passed through a UV spectrophotometer to get the absorbance profile at 254nm. The individual fractions were subjected to Western blotting to probe for ribosomal protein RPLP0 and RNA isolation/qPCR to probe for candidate mRNAs.

To identify actively translating polysomes, the cells were treated with 1mM Puromycin (P8833, Sigma) instead of Cycloheximide. The Puromycin-sensitive fractions 7-11 were considered as the actively translating fractions whereas the Puromycin-insensitive fractions 1-6 were considered as the non-translating pool. The ratio was considered as shown – *Ratio of (Translating pool / Non-translating pool) = Ratio of (sum of percentage of mRNA or protein in fractions 7-11 / sum of percentage of mRNA or protein in fractions 1-6)*

To further dissect the distribution of mRNAs/proteins in the non-translating pool, fractions 1-6 were further divided into fractions 1-3 (inhibitory complex) and fractions 4-6 (ribosomal subunits and monosomes) *. The enrichment of the mRNAs/proteins in the inhibitory complex was calculated as – *Sum of percentage of mRNAs or proteins in fractions 1-3 / Sum of percentage of mRNAs or proteins in fraction 4-6*

*In this case, the percentage was calculated with respect to distribution in fractions 1-6*

### Calcium imaging

Calcium imaging was done on DIV15 neurons plated on Nunc glass-bottomed imaging dishes. The imaging and washes were performed with ACSF media (120mM NaCl, 3mM KCl, 1mM MgCl_2_, 3mM NaHCO_3_, 1.25mM NaH_2_PO_4_, 15mM HEPES, 2mM CaCl_2_, 30mM glucose, pH 7.4). The cells were washed once with ACSF and incubated at 37°C with 1ml of freshly prepared Fluo4-AM dye solution (1μM Fluo4-AM and 0.002% Pluronic acid in ACSF) (F14217, ThermoFisher Scientific) for 20 minutes. They were given two washes and incubated in ACSF at 37°C for 10-20minutes before imaging. The drugs Nifedipine (50μM) and MK801 (25μM) were added during this incubation step. The neurons were imaged using Olympus FV300 confocal laser scanning inverted microscope with 20X objective, illuminated with 488nm lasers. The neurons were imaged for a total time of 7 minutes at a rate of 3 seconds per frame (140 frames in total). They were imaged in basal condition for 1 minute (20 frames). Followed by that, they were imaged for 5 minutes (100 frames) on APOE recombinant protein (20nM) addition. Finally, they were imaged for 1 minute (20 frames) on the addition of Ionomycin solution (10μM Ionomycin with 10mM CaCl_2_) (407950, Sigma). The images obtained were analyzed using the Time-Series Analyzer plug-in on FIJI. The average intensities were obtained for the selected ROIs (ROIs were drawn manually and the Ionomycin responsive cells were included). The change in fluorescent intensity at each frame was normalized to the initial fluorescent intensity of the first frame (F0) for each ROI. The normalized change in fluorescent intensity (ΔF/F0) was plotted along the time axis and used for statistical analysis as well.

### Differentiation of iPSCs to neurons

The protocol for neural differentiation was adapted from Yichen Shi *et al* (2012)^47^ and Yu Zhang *et al* (2017)^46^ with some modifications. In brief, iPSCs were expanded in mTeSR until 70-80% confluency was reached. The Neural Basic Media (NBM) for differentiation contained 50% DMEM F-12 (21331–020, ThermoFisher Scientific), 50% Neurobasal, 0.1% PenStrep, Glutamax, N2 (17502–048, ThermoFisher Scientific), and B27 without Vitamin A (12587–010, ThermoFisher Scientific). The iPSCs were subjected to monolayer Dual SMAD inhibition by changing the media to Neural Induction Media (NIM) which is composed of NBM supplemented with small molecules SB431542 (10μM, an inhibitor of TGFβ pathway) (72232, Stem Cell Technologies) and LDN193189 (0.1μM, an inhibitor of BMP pathway) (72142, Stem Cell Technologies). The cells were subjected to neural induction for 12-15 days by changing the NIM every day. After the induction, the monolayer was dissociated using Accutase (A6964, Sigma) and the cells were plated in NIM containing 10μM ROCK inhibitor (Y0503, Sigma) overnight on pre-coated poly-L-ornithine/laminin dishes. Poly-L-Ornithine (1:10 dilution) (P4957, Sigma) followed by Laminin (5μg/ml) (L2020, Sigma) coating was used for maintenance of neural progenitors and their terminal differentiation. Expansion of the neural progenitor cells was carried out in Neural Expansion Media (NEM) which is composed of NBM supplemented with FGF (10ng/ml) (100-18C, Peprotech) and EGF (10ng/ml) (AF-100-15, Pepotech). Neuronal maturation and terminal differentiation were achieved by plating the neural stem cells in the density of 25,000-35,000 cells/cm^2^ in the Neural Maturation Media (NMM) composed of NBM supplemented with BDNF (20ng/ml) (450-02, Peprotech), GDNF (10ng/ml) (450-10, Peprotech), L-Ascorbic Acid (200μM) (A4403, Sigma) and db-Camp (50μM) (D0627, Sigma). The neurons were subjected to maturation for a period of 4-5 weeks by supplementing them with NMM after every 4-5 days.

### Statistical analysis

All statistical analyses were performed using Graph Pad Prism software. The normality of the data was checked using the Kolmogorov-Smirnov test. For experiments with less than 5 data points, parametric statistical tests were applied. Data were represented as mean ± SEM in all biochemical experiment graphs. FUNCAT imaging data was represented as boxes and whiskers with all the individual data points. Statistical significance was calculated using Unpaired Student’s t-test (2 tailed with equal variance) in cases where 2 groups were being compared. One-way ANOVA was used for multiple group comparisons, followed by Tukey’s multiple comparison test. P-value less than 0.05 was considered to be statistically significant.

### Antibodies used for Western blotting

Antibodies used for Western blotting –
| Protein | Dilution | Catalog number, Company |
| --- | --- | --- |
| eEF2 | 1:1000 | 2332S, Cell Signaling Technologies |
| p-eEF2 | 1:2000 | 2331S, Cell Signaling Technologies |
| ERK | 1:1000 | 9102, Cell Signaling Technologies |
| p-ERK | 1:1000 | 9101, Cell Signaling Technologies |
| APOE | 1:1000 | NB110-60531, Novus Biologicals |
| Tuj1 | 1:4000 | T8578, Sigma |
| RPLP0 | 1:4000 | ab101279, Abcam |
| RPS6 | 1:3000 | 2217, Cell Signaling Technologies |
| PTEN | 1:1000 | 9552S, Cell Signaling Technologies |
| PSD95 | 1:1000 | P246, Sigma |
| α-tubulin | 1:5000 | T9026, Sigma |
| eIF2 | 1:1000 | 9722S, Cell Signaling Technologies |
| p-eIF2 | 1:1000 | 9721S, Cell Signaling Technologies |
| Secondary Rabbit HRP | 1:5000 | A0545, Sigma |
| Secondary Mouse HRP | 1:5000 | 31430, Thermofisher Scientific |

### Antibodies used for Immunostaining

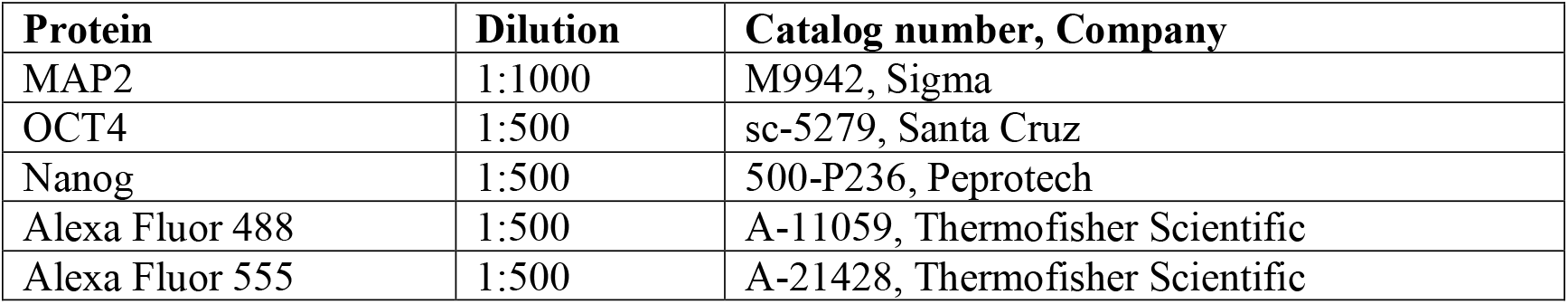

Antibodies used for Immunostaining –
| Protein | Dilution | Catalog number, Company |
| --- | --- | --- |
| MAP2 | 1:1000 | M9942, Sigma |
| OCT4 | 1:500 | sc-5279, Santa Cruz |
| Nanog | 1:500 | 500-P236, Peprotech |
| Alexa Fluor 488 | 1:500 | A-11059, Thermofisher Scientific |
| Alexa Fluor 555 | 1:500 | A-21428, Thermofisher Scientific |

### Primers used for RT-PCR

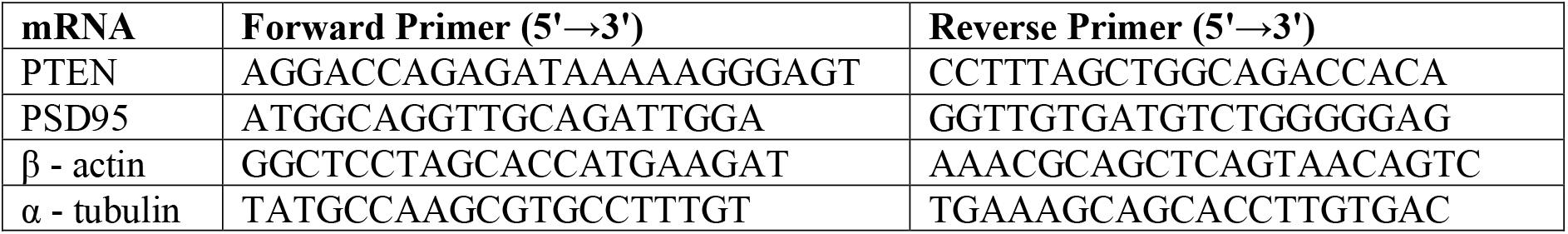

## Supporting information

Supplementary figures with legends

## Acknowledgments

We are thankful to all the central facilities at NCBS-inStem, especially the Central Imaging and Flow Cytometry Facility (CIFF), Animal House Facility, and Stem Cell Facility. We thank our collaborators and colleagues for their invaluable suggestions and discussions.

## Author Contributions

SR and RM designed and conceptualized the experiments. SR performed the biochemistry and imaging experiments with rat neurons, human neurons, and synaptoneurosomes. VJ performed the polysome profiling assays with NMDAR stimulation and 1-minute APOE treatment. SC performed the calcium imaging for chronic (24 hours) APOE treatment. SK performed the experiments with mouse astrocyte conditioned media and mouse neurons. GG provided the APOE animal resources and critical suggestions for the project. KF provided the APOE iPSCs and contributed towards the manuscript with important suggestions. BC and BH generated and provided the APOE iPSC lines. The manuscript was written by SR and RM.

## References

1. Hardy, J. A Hundred Years of Alzheimer’s Disease Research. Neuron vol. 52 (2006).

2. Shankar, G. M. & Walsh, D. M. Alzheimer’s disease: Synaptic dysfunction and Aβ. Molecular Neurodegeneration 4, (2009).

3. Sutton, M. A. & Schuman, E. M. Dendritic Protein Synthesis, Synaptic Plasticity, and Memory. Cell (2006) doi:10.1016/j.cell.2006.09.014.

4. Kelleher, R. J., Govindarajan, A. & Tonegawa, S. Translational regulatory mechanisms in persistent forms of synaptic plasticity. Neuron vol. 44 (2004).

5. Suzanne Zukin, R., Richter, J. D. & Bagni, C. Signals, synapses, and synthesis: How new proteins control plasticity. Frontiers in Neural Circuits vol. 3 (2009).

6. Chen, Y. C., Chang, Y. W. & Huang, Y. S. Dysregulated Translation in Neurodevelopmental Disorders: An Overview of Autism-Risk Genes Involved in Translation. Developmental Neurobiology vol. 79 (2019).

7. Ma, T. et al. Dysregulation of the mTOR pathway mediates impairment of synaptic plasticity in a mouse model of Alzheimer’s disease. PLoS ONE 5, (2010).

8. Yang, W. et al. Repression of the eIF2α kinase PERK alleviates mGluR-LTD impairments in a mouse model of Alzheimer’s disease. Neurobiology of Aging 41, (2016).

9. Beckelman, B. C. et al. Dysregulation of Elongation Factor 1A Expression is Correlated with Synaptic Plasticity Impairments in Alzheimer’s Disease. Journal of Alzheimer’s Disease 54, (2016).

10. An, W. L. et al. Up-regulation of phosphorylated/activated p70 S6 kinase and its relationship to neurofibrillary pathology in Alzheimer’s disease. American Journal of Pathology 163, (2003).

11. Li, X. et al. Phosphorylated eukaryotic translation factor 4E is elevated in Alzheimer brain. NeuroReport 15, (2004).

12. Ding, Q. Ribosome Dysfunction Is an Early Event in Alzheimer’s Disease. Journal of Neuroscience 25, 9171–9175 (2005).

13. Ding, Q., Markesbery, W. R., Cecarini, V. & Keller, J. N. Decreased RNA, and increased RNA oxidation, in ribosomes from early Alzheimer’s disease. Neurochemical Research 31, 705–710 (2006).

14. Hernández-Ortega, K., Garcia-Esparcia, P., Gil, L., Lucas, J. J. & Ferrer, I. Altered Machinery of Protein Synthesis in Alzheimer’s: From the Nucleolus to the Ribosome. Brain Pathology 26, 593–605 (2016).

15. Elder, M. K. et al. Dysregulation of the de novo proteome accompanies pathology progression in the APP/PS1 mouse model. PrePrint (2020).

16. Cefaliello, C. et al. Deregulated Local Protein Synthesis in the Brain Synaptosomes of a Mouse Model for Alzheimer’s Disease. Molecular Neurobiology 57, (2020).

17. Meier, S. et al. Pathological Tau Promotes Neuronal Damage by Impairing Ribosomal Function and Decreasing Protein Synthesis. Journal of Neuroscience 36, 1001–1007 (2016).

18. Evans, H. T., Benetatos, J., van Roijen, M., Bodea, L. & Götz, J. Decreased synthesis of ribosomal proteins in tauopathy revealed by non-canonical amino acid labelling. The EMBO Journal 38, (2019).

19. Ahmad, F. et al. Reactive Oxygen Species-Mediated Loss of Synaptic Akt1 Signaling Leads to Deficient Activity-Dependent Protein Translation Early in Alzheimer’s Disease. Antioxidants and Redox Signaling 27, (2017).

20. Tanzi, R. E. The genetics of Alzheimer disease. Cold Spring Harbor Perspectives in Medicine 2, (2012).

21. Kim, J., Basak, J. M. & Holtzman, D. M. The Role of Apolipoprotein E in Alzheimer’s Disease. Neuron 63, 287–303 (2009).

22. Liu, C. C., Kanekiyo, T., Xu, H. & Bu, G. Apolipoprotein e and Alzheimer disease: Risk, mechanisms and therapy. Nature Reviews Neurology vol. 9 (2013).

23. Rodriguez, G. A., Burns, M. P., Weeber, E. J. & Rebeck, G. W. Young APOE4 targeted replacement mice exhibit poor spatial learning and memory, with reduced dendritic spine density in the medial entorhinal cortex. Learning and Memory 20, (2013).

24. Yong, S. M., Lim, M. L., Low, C. M. & Wong, B. S. Reduced neuronal signaling in the ageing apolipoprotein-E4 targeted replacement female mice. Scientific Reports 4, (2014).

25. Teter, B. et al. Defective neuronal sprouting by human apolipoprotein E4 is a gain-of-negative function. Journal of Neuroscience Research 68, (2002).

26. Nathan, B. P. et al. Differential effects of apolipoproteins E3 and E4 on neuronal growth in vitro. Science 264, (1994).

27. Dumanis, S. B. et al. ApoE4 decreases spine density and dendritic complexity in cortical neurons in vivo. Journal of Neuroscience 29, (2009).

28. Reas, E. T. et al. Effects of APOE on cognitive aging in community-dwelling older adults. Neuropsychology 33, (2019).

29. de Jager, P. L. et al. A genome-wide scan for common variants affecting the rate of age-related cognitive decline. Neurobiology of Aging 33, (2012).

30. Wisdom, N. M., Callahan, J. L. & Hawkins, K. A. The effects of apolipoprotein E on non-impaired cognitive functioning: A meta-analysis. Neurobiology of Aging 32, (2011).

31. Small, B. J., Rosnick, C. B., Fratiglioni, L. & Bäckman, L. Apolipoprotein E and cognitive performance: A meta-analysis. Psychology and Aging 19, (2004).

32. Chen, Y., Durakoglugil, M. S., Xian, X. & Herz, J. ApoE4 reduces glutamate receptor function and synaptic plasticity by selectively impairing ApoE receptor recycling. Proceedings of the National Academy of Sciences of the United States of America 107, (2010).

33. Bacskai, B. J., Xia, M. Q., Strickland, D. K., Rebeck, G. W. & Hyman, B. T. The endocytic receptor protein LRP also mediates neuronal calcium signaling via N-methyl-D-aspartate receptors. Proceedings of the National Academy of Sciences of the United States of America 97, (2000).

34. Sheng, Z., Prorok, M., Brown, B. E. & Castellino, F. J. N-methyl-d-aspartate receptor inhibition by an apolipoprotein E-derived peptide relies on low-density lipoprotein receptor-associated protein. Neuropharmacology 55, (2008).

35. Nakajima, C. et al. Low density lipoprotein receptor-related protein 1 (LRP1) modulates N-methyl-D-aspartate (NMDA) receptor-dependent intracellular signaling and NMDA-induced regulation of postsynaptic protein complexes. Journal of Biological Chemistry 288, (2013).

36. May, P. et al. Neuronal LRP1 Functionally Associates with Postsynaptic Proteins and Is Required for Normal Motor Function in Mice. Molecular and Cellular Biology 24, (2004).

37. Huang, Y. W. A., Zhou, B., Wernig, M. & Südhof, T. C. ApoE2, ApoE3, and ApoE4 Differentially Stimulate APP Transcription and Aβ Secretion. Cell 168, 427–441.e21 (2017).

38. Huang, Y. W. A., Zhou, B., Nabet, A. M., Wernig, M. & Südhof, T. C. Differential Signaling Mediated by ApoE2, ApoE3, and ApoE4 in Human Neurons Parallels Alzheimer’s Disease Risk. Journal of Neuroscience 39, 7408–7427 (2019).

39. Hoe, H. S., Harris, D. C. & Rebeck, G. W. Multiple pathways of apolipoprotein E signaling in primary neurons. Journal of Neurochemistry 93, (2005).

40. Wang, C. et al. Human apoE4-targeted replacement mice display synaptic deficits in the absence of neuropathology. Neurobiology of Disease 18, 390–398 (2005).

41. Liu, D. S. et al. APOE4 enhances age-dependent decline in cognitive function by down-regulating an NMDA receptor pathway in EFAD-Tg mice. Molecular Neurodegeneration 10, 1–17 (2015).

42. Schmid, B. et al. Generation of a set of isogenic, gene-edited iPSC lines homozygous for all main APOE variants and an APOE knock-out line. Stem Cell Research (2019) doi:10.1016/j.scr.2018.11.010.

43. Heise, C. et al. Elongation factor-2 phosphorylation in dendrites and the regulation of dendritic mRNA translation in neurons. Frontiers in Cellular Neuroscience vol. 8 (2014).

44. Ghosh Dastidar, S. et al. Distinct regulation of bioenergetics and translation by group I mGluR and NMDAR. EMBO reports (2020) doi:10.15252/embr.201948037.

45. Dieck, S. T. et al. Metabolic labeling with noncanonical amino acids and visualization by chemoselective fluorescent tagging. Current Protocols in Cell Biology 1, (2012).

46. Zhang, Y. et al. Patient iPSC-Derived Neurons for Disease Modeling of Frontotemporal Dementia with Mutation in CHMP2B. Stem Cell Reports 8, (2017).

47. Shi, Y., Kirwan, P. & Livesey, F. J. Directed differentiation of human pluripotent stem cells to cerebral cortex neurons and neural networks. Nature Protocols 7, 1836–1846 (2012).

48. Qiu, Z., Crutcher, K. A., Hyman, B. T. & Rebeck, G. W. apoE isoforms affect neuronal N-methyl-D-aspartate calcium responses and toxicity via receptor-mediated processes. Neuroscience 122, 291–303 (2003).

49. Kute, P. M., Ramakrishna, S., Neelagandan, N., Chattarji, S. & Muddashetty, Ravi. S. NMDAR mediated translation at the synapse is regulated by MOV10 and FMRP. Molecular Brain 12, 65 (2019).

50. Kuzniewska, B. et al. Mitochondrial protein biogenesis in the synapse is supported by local translation. EMBO reports 21, (2020).

51. Pakos-Zebrucka, K. et al. The integrated stress response. EMBO reports 17, (2016).

52. Dittmer, P. J., Wild, A. R., Dell’Acqua, M. L. & Sather, W. A. STIM1 Ca2+ Sensor Control of L-type Ca2+-Channel-Dependent Dendritic Spine Structural Plasticity and Nuclear Signaling. Cell Reports 19, 321–334 (2017).

53. Wang, C. et al. Gain of toxic apolipoprotein E4 effects in human iPSC-derived neurons is ameliorated by a small-molecule structure corrector article. Nature Medicine 24, (2018).

54. Ulrich, J. D. et al. In vivo measurement of apolipoprotein e from the brain interstitial fluid using microdialysis. Molecular Neurodegeneration 8, (2013).

55. Scheetz, A. J., Nairn, A. C. & Constantine-Paton, M. NMDA receptor-mediated control of protein synthesis at developing synapses. Nature Neuroscience (2000) doi:10.1038/72915.

56. Hoeffer, C. A. & Klann, E. NMDA receptors and translational control. in Biology of the NMDA Receptor (2008). doi:10.1201/9781420044157.ch6.

57. Higley, M. J. & Sabatini, B. L. Calcium signaling in dendritic spines. Cold Spring Harbor Perspectives in Biology 4, (2012).

58. Lau, C. G. et al. Regulation of NMDA receptor Ca2+ signalling and synaptic plasticity. Biochemical Society Transactions 37, (2009).

59. Papadia, S. & Hardingham, G. E. The Dichotomy of NMDA Receptor Signaling. The Neuroscientist 13, (2007).

60. Hiester, B. G. et al. L-Type Voltage-Gated Ca2+ Channels Regulate Synaptic-Activity-Triggered Recycling Endosome Fusion in Neuronal Dendrites. Cell Reports 21, (2017).

61. Emptage, N., Bliss, T. V. P. & Fine, A. Single synaptic events evoke NMDA receptor-mediated release of calcium from internal stores in hippocampal dendritic spines. Neuron 22, (1999).

62. Lee, K. F. H., Soares, C., Thivierge, J. P. & Béïque, J. C. Correlated Synaptic Inputs Drive Dendritic Calcium Amplification and Cooperative Plasticity during Clustered Synapse Development. Neuron 89, (2016).

63. Griffith, T., Tsaneva-Atanasova, K. & Mellor, J. R. Control of Ca2+ Influx and Calmodulin Activation by SK-Channels in Dendritic Spines. PLoS Computational Biology 12, (2016).

64. Rajadhyaksha, A. et al. L-type Ca2+ channels are essential for glutamate-mediated CREB phosphorylation and c-fos gene expression in striatal neurons. Journal of Neuroscience 19, (1999).

65. Colbran, R. J. Protein phosphatases and calcium/calmodulin-dependent protein kinase II-dependent synaptic plasticity. Journal of Neuroscience vol. 24 (2004).

66. Tolar, M. et al. Truncated apolipoprotein E (ApoE) causes increased intracellular calcium and may mediate ApoE neurotoxicity. Journal of Neuroscience 19, (1999).

67. Veinbergs, I., Everson, A., Sagara, Y. & Masliah, E. Neurotoxic effects of apolipoprotein E4 are mediated via dysregulation of calcium homeostasis. Journal of Neuroscience Research 67, (2002).

68. Xu, D. & Peng, Y. Apolipoprotein E 4 triggers multiple pathway-mediated Ca2+ overload, causes CaMK II phosphorylation abnormity and aggravates oxidative stress caused cerebral cortical neuron damage. European review for medical and pharmacological sciences 21, (2017).

69. Qiu, Z., Hyman, B. T. & Rebeck, G. W. Apolipoprotein E receptors mediate neurite outgrowth through activation of p44/42 mitogen-activated protein kinase in primary neurons. Journal of Biological Chemistry 279, 34948–34956 (2004).

70. Ohkubo, N. et al. Apolipoprotein E4 Stimulates cAMP Response Element-binding Protein Transcriptional Activity through the Extracellular Signal-regulated Kinase Pathway. Journal of Biological Chemistry 276, 3046–3053 (2001).

71. Chen, Y. et al. Reelin modulates NMDA receptor activity in cortical neurons. Journal of Neuroscience 25, (2005).

72. Lane-Donovan, C. & Herz, J. ApoE, ApoE Receptors, and the Synapse in Alzheimer’s Disease. Trends in Endocrinology and Metabolism 28, 273–284 (2017).

73. Lin, Y. T. et al. APOE4 Causes Widespread Molecular and Cellular Alterations Associated with Alzheimer’s Disease Phenotypes in Human iPSC-Derived Brain Cell Types. Neuron 98, 1141–1154.e7 (2018).

74. Texidó, L., Martín-Satué, M., Alberdi, E., Solsona, C. & Matute, C. Amyloid β peptide oligomers directly activate NMDA receptors. Cell Calcium 49, (2011).

75. Ravindran, S., Nalavadi, V. C. & Muddashetty, R. S. BDNF induced translation of limk1 in developing neurons regulates dendrite growth by fine-tuning cofilin1 activity. Frontiers in Molecular Neuroscience (2019) doi:10.3389/fnmol.2019.00064.

76. Muddashetty, R. S., Kelić, S., Gross, C., Xu, M. & Bassell, G. J. Dysregulated metabotropic glutamate receptor-dependent translation of AMPA receptor and postsynaptic density-95 mRNAs at synapses in a mouse model of fragile X syndrome. Journal of Neuroscience (2007) doi:10.1523/JNEUROSCI.0937-07.2007.

77. Paul, A. et al. Differential regulation of syngap1 translation by FMRP modulates eEF2 mediated response on NMDAR activity. Frontiers in Molecular Neuroscience (2019) doi:10.3389/fnmol.2019.00097.

78. Takahashi, R. H. et al. Oligomerization of Alzheimer’s β-Amyloid within Processes and Synapses of Cultured Neurons and Brain. Journal of Neuroscience 24, (2004).

79. Piedrahita, J. A., Zhang, S. H., Hagaman, J. R., Oliver, P. M. & Maeda, N. Generation of mice carrying a mutant apolipoprotein E gene inactivated by gene targeting in embryonic stem cells. Proceedings of the National Academy of Sciences of the United States of America 89, (1992).

