## Supplementary figures with legends for "APOE4 affects basal and NMDAR mediated protein synthesis in neurons by perturbing calcium homeostasis"

### Supplementary Figure 1

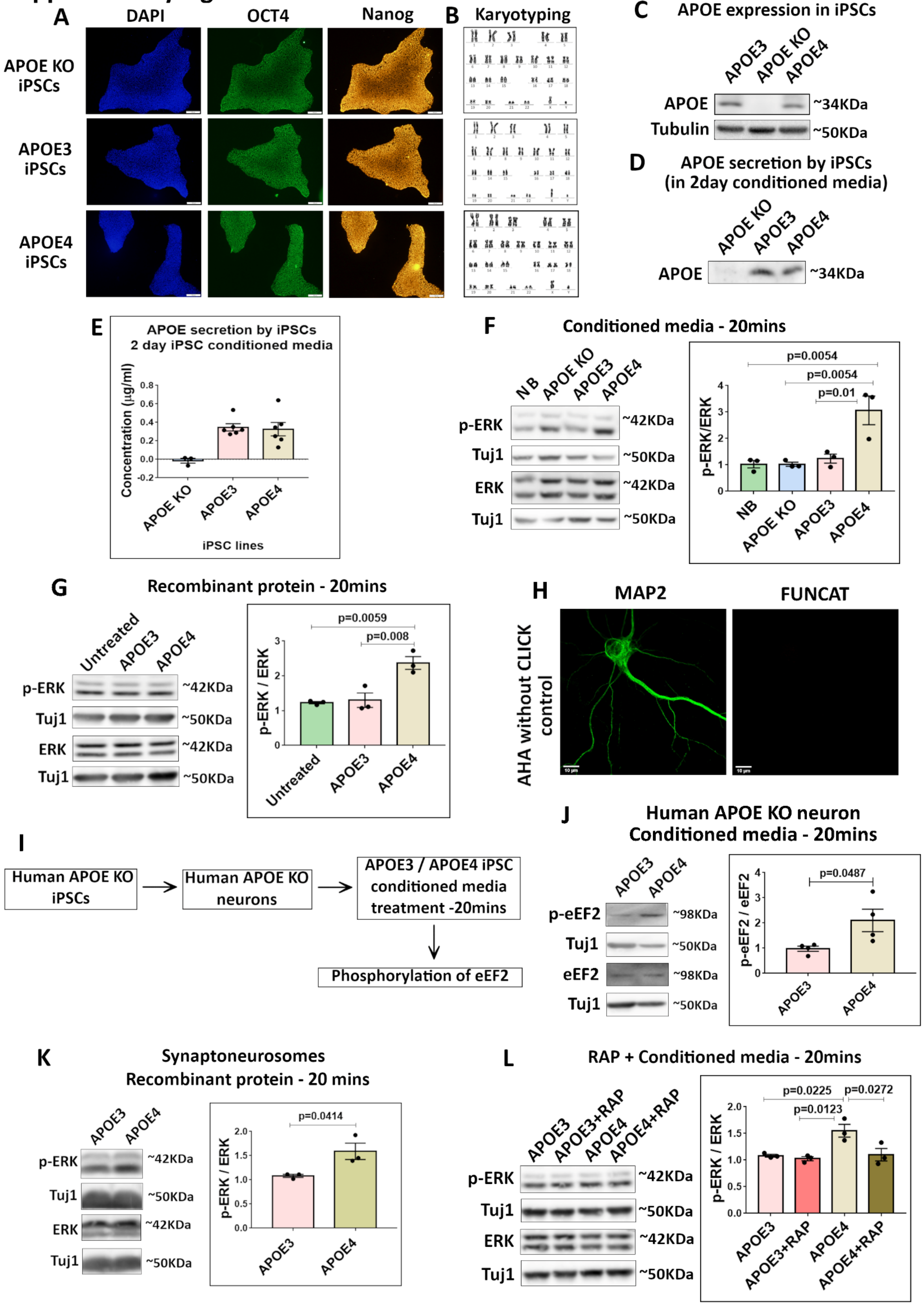

**Supplementary Figure 1 -**

**A** – APOE KO, APOE3 and APOE4 iPSCs were characterized for the expression of nuclear marker DAPI (left) and pluripotency markers OCT4 (middle) and NANOG (right). The representative images for the same are shown (Scale bar - 100µM)

**B** – The karyotyping profile of APOE KO, APOE3 and APOE4 iPSCs

**C** - APOE KO, APOE3 and APOE4 iPSC cell lysates were probed for the expression of APOE. Representative immunoblots indicating the levels of APOE and loading control Tubulin.

**D** - APOE KO, APOE3 and APOE4 iPSCs (once reached 50% confluency) were maintained in neurobasal for 48 hours. Representative immunoblots indicate the APOE secreted by the iPSCs in 2-day conditioned neurobasal media.

**E** – The amount of APOE secreted by the iPSCs in 2-day conditioned neurobasal media was estimated through ELISA. The graph indicates the average APOE concentration (µg/ml) in 2-day iPSC conditioned media.

**F** - Rat primary cortical neurons (DIV15) were treated with APOE (10-15nM) from iPSC conditioned media for 20 minutes and probed for phosphorylation of ERK. Left - representative immunoblots indicating levels of phospho-ERK, ERK and Tuj1; Right - graph indicating ratio of phospho-ERK to ERK normalized to Tuj1. Data is represented as mean +/- SEM. N=3, One-way ANOVA (p=0.0031) followed by Tukey's multiple comparison test.

**G** - Rat primary cortical neurons (DIV15) were treated with recombinant APOE protein (15nM) for 20minutes and probed for phosphorylation of ERK. Left - representative immunoblots indicating levels of phospho-ERK, ERK and Tuj1; Right - graph indicating ratio of phospho-ERK to ERK normalized to Tuj1. Data is represented as mean +/- SEM. N=3, One-way ANOVA (p=0.0041) followed by Tukey's multiple comparison test.

**H** - Rat primary cortical neurons (DIV15) were subjected to fluorescent non-canonical amino acid tagging (FUNCAT) along with immunostaining for MAP2. The representative images for MAP2 and FUNCAT fluorescent signal under the control condition of AHA without Click reaction is shown.

**I** – Experimental workflow. Human APOE KO neurons were derived from human APOE KO iPSCs and subjected to APOE3 or APOE4 iPSC conditioned media treatment for 20 minutes. They were probed for phosphorylation of eEF2 as a readout for global protein synthesis.

**J** – Human APOE KO neurons (4 weeks into neuronal maturation) were treated with APOE3/APOE4 (10-15nM) iPSC conditioned media for 20 minutes and probed for phosphorylation of eEF2. Left - representative immunoblots indicating levels of phospho-eEF2, eEF2 and Tuj1; Right - graph indicating ratio of phospho-eEF2 to eEF2 normalized to Tuj1. Data is represented as mean +/- SEM. N=4, Unpaired Student's t-test.

**K** - Synaptoneurosomes prepared from P30 rat cortices were treated with recombinant APOE protein (15nM) for 20 minutes and probed for phosphorylation of ERK. Left - representative immunoblots indicating levels of phospho-ERK, ERK and Tuj1; Right - graph indicating ratio of phospho-ERK to ERK normalized to Tuj1. Data is represented as mean +/- SEM. N=3, Unpaired Student's t-test.

**L** - Rat primary cortical neurons (DIV15) were treated with APOE receptor antagonist RAP (200nM) along with APOE (10-15nM) from iPSC conditioned media for 20 minutes and probed for phosphorylation of ERK. Left - representative immunoblots indicating levels of phospho-ERK, ERK and Tuj1; Right - graph indicating ratio of phospho-ERK to ERK normalized to Tuj1. Data is represented as mean +/- SEM. N=3, One-way ANOVA (p=0.0097) followed by Tukey's multiple comparison test.

#### Supplementary Figure 2

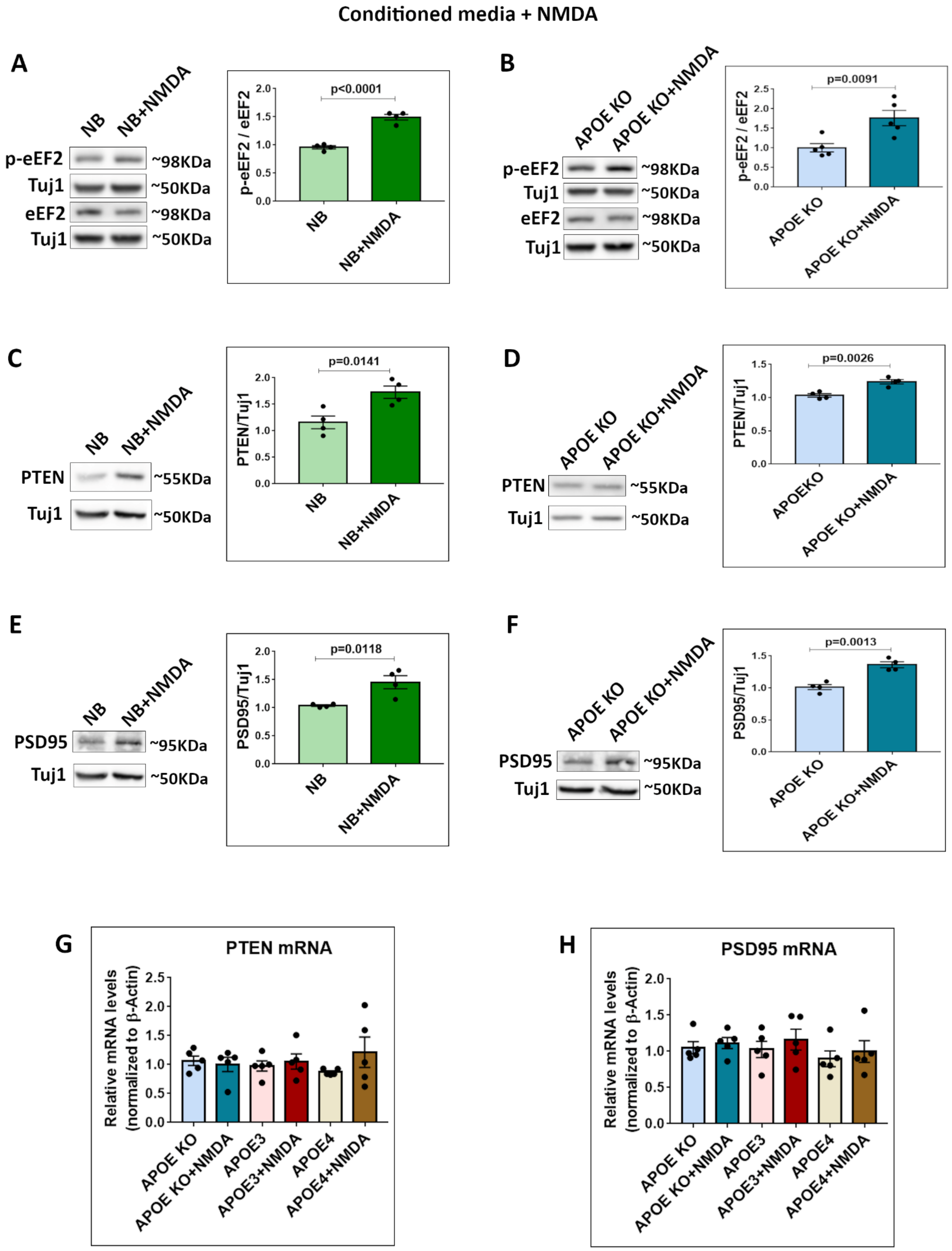

**Supplementary Figure 2 -**

**A** - Rat primary cortical neurons (DIV15) were treated with neurobasal media for 20 minutes along with NMDAR stimulation for 5 minutes (20µM NMDA). The cell lysates were probed for phosphorylation of eEF2. Left - representative immunoblots indicating levels of phospho-eEF2, eEF2 and Tuj1; Right - graph indicating ratio of phospho-eEF2 to eEF2 normalized to Tuj1. Data is represented as mean +/- SEM. N=4, Unpaired Student's t-test.

**B** - Rat primary cortical neurons (DIV15) were treated with APOE KO conditioned media for 20 minutes along with NMDAR stimulation for 5 minutes (20µM NMDA). The cell lysates were probed for phosphorylation of eEF2. Left - representative immunoblots indicating levels of phospho-eEF2, eEF2 and Tuj1; Right - graph indicating ratio of phospho-eEF2 to eEF2 normalized to Tuj1. Data is represented as mean +/- SEM. N=5, Unpaired Student's t-test.

**C** - Rat primary cortical neurons (DIV15) were treated with neurobasal media for 20 minutes along with NMDAR stimulation for 5 minutes (20µM NMDA). The cell lysates were probed for PTEN protein. Left - representative immunoblots indicating levels of PTEN and Tuj1; Right - graph indicating PTEN levels normalized to Tuj1. Data is represented as mean +/- SEM. N=4, Unpaired Student's t-test.

**D** - Rat primary cortical neurons (DIV15) were treated with APOE KO conditioned media for 20 minutes along with NMDAR stimulation for 5 minutes (20µM NMDA). The cell lysates were probed for PTEN protein. Left - representative immunoblots indicating levels of PTEN and Tuj1; Right - graph indicating PTEN levels normalized to Tuj1. Data is represented as mean +/- SEM. N=4, Unpaired Student's t-test.

**E** - Rat primary cortical neurons (DIV15) were treated with neurobasal media for 20 minutes along with NMDAR stimulation for 5 minutes (20µM NMDA). The cell lysates were probed for PSD95 protein. Left - representative immunoblots indicating levels of PSD95 and Tuj1; Right - graph indicating PSD95 levels normalized to Tuj1. Data is represented as mean +/- SEM. N=4, Unpaired Student's t-test.

**F** - Rat primary cortical neurons (DIV15) were treated with APOE KO conditioned media for 20 minutes along with NMDAR stimulation for 5 minutes (20µM NMDA). The cell lysates were probed for PSD95 protein. Left - representative immunoblots indicating levels of PSD95 and Tuj1; Right - graph indicating PSD95 levels normalized to Tuj1. Data is represented as mean +/- SEM. N=4, Unpaired Student's t-test.

**G** - Rat primary cortical neurons (DIV15) were treated with APOE KO/APOE3/APOE4 (10-15nM) conditioned media for 20minutes along with NMDAR stimulation for 5minutes (20µM NMDA) and subjected to RT-PCR to measure the levels of PTEN mRNA. The graph indicates the copy number of PTEN mRNA normalized to copy number of β-actin mRNA under different treatment conditions. Data is represented as mean +/- SEM, N=5. One-way ANOVA (ns).

**H** - Rat primary cortical neurons (DIV15) were treated with APOE KO/APOE3/APOE4 (10-15nM) conditioned media for 20minutes along with NMDAR stimulation for 5minutes (20µM NMDA) and subjected to RT-PCR to measure the levels of PSD95 mRNA. The graph indicates the copy number of PSD95 mRNA normalized to copy number of β-actin mRNA under different treatment conditions. Data is represented as mean +/- SEM, N=5. One-way ANOVA (ns).

#### Supplementary Figure 3

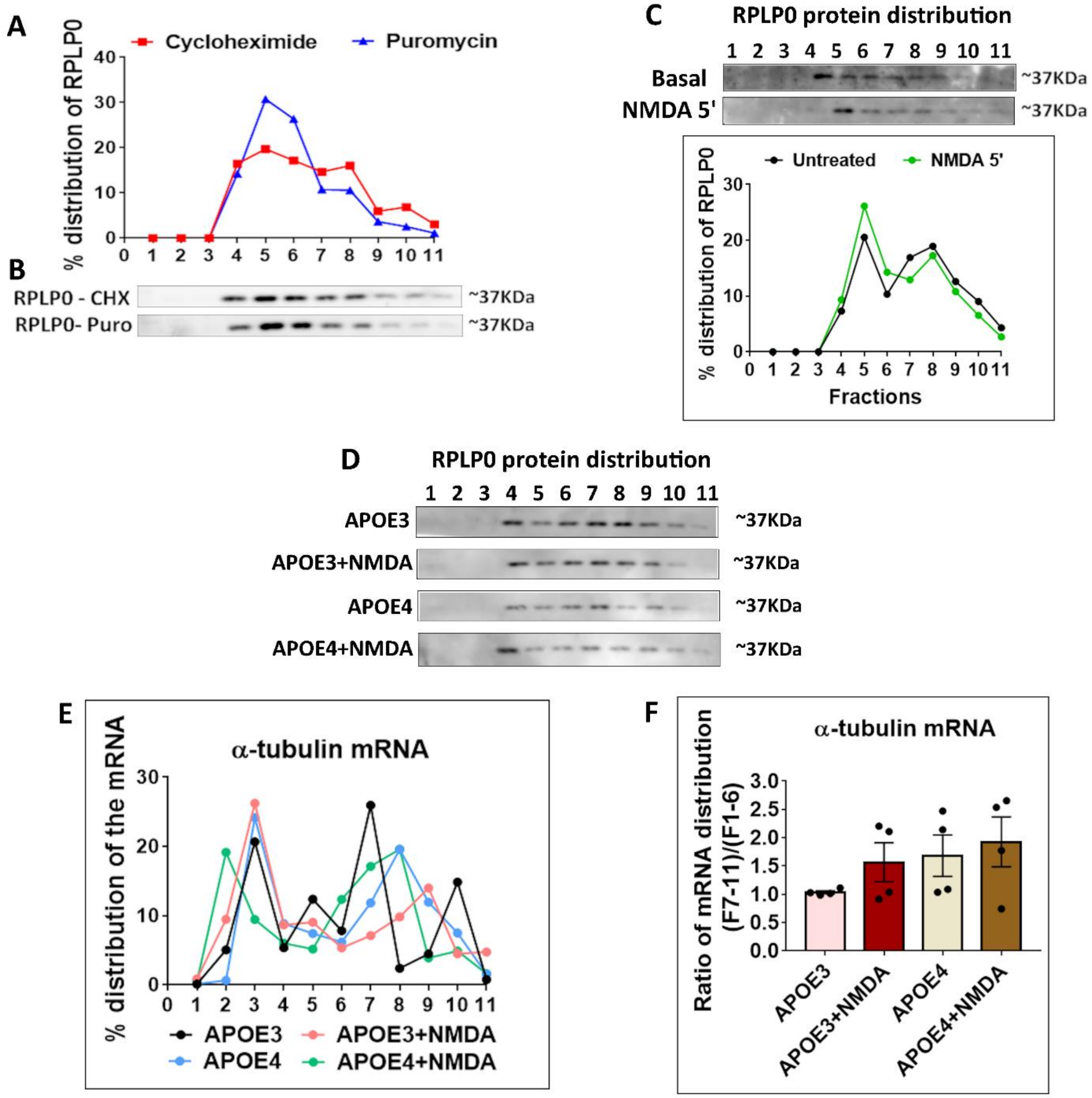

**Supplementary Figure 3 –**

**A -** Rat primary cortical neurons (DIV15) were subjected to polysome profiling after treatment with Cycloheximide (CHX) and Puromycin (Puro)*. The distribution of ribosomal RPLP0 on the linear sucrose gradient was quantified using immunoblots shown in Fig S3B. The graph represents the percentage distribution of RPLP0 in each fraction under Cycloheximide (CHX) and Puromycin (Puro) treatment conditions.

*Puromycin disrupts actively translating polysomes and hence helps in identifying this pool.

**B** – The immunoblots showing the distribution of ribosomal protein RPLP0 in the individual fractions on the linear sucrose gradient under Cycloheximide and Puromycin treatment conditions.

**C -** Rat primary cortical neurons (DIV15) were subjected to NMDAR stimulation for 5 minutes (20µM NMDA). The cell lysates were subjected to polysome profiling and probed for the distribution of ribosomal protein RPLP0. Top - immunoblots indicating the distribution of RPLP0 in individual fractions; bottom - line graph indicating the percentage distribution of RPLP0 in individual fractions under basal and 5-minute NMDA stimulated conditions.

**D -** Rat primary cortical neurons (DIV15) were treated with APOE3/APOE4 (10-15nM) conditioned media for 20 minutes along with NMDAR stimulation for 5minutes (20µM NMDA). The cell lysates were subjected to polysome profiling and probed for ribosomal protein RPLP0. The immunoblots show the distribution of RPLP0 in individual fractions of the linear sucrose gradient under different conditions.

Rat primary cortical neurons (DIV15) were treated with APOE3/APOE4 (10-15nM) conditioned media for 20 minutes along with NMDAR stimulation for 5 minutes (20µM NMDA). The cell lysates were subjected to polysome profiling and probed for the distribution of the following candidate mRNAs in each fraction using RT-PCR method –

**E -** The line graph represents the percentage distribution of α-tubulin mRNA in each fraction under different treatment conditions.

**F -** The graph represents the ratio of α-tubulin mRNA distribution in fractions 7-11 (translating pool) to fractions 1-6 (non-translating pool). Data is represented as mean +/- SEM. N=3, One-way ANOVA (ns).

#### Supplementary Figure 4

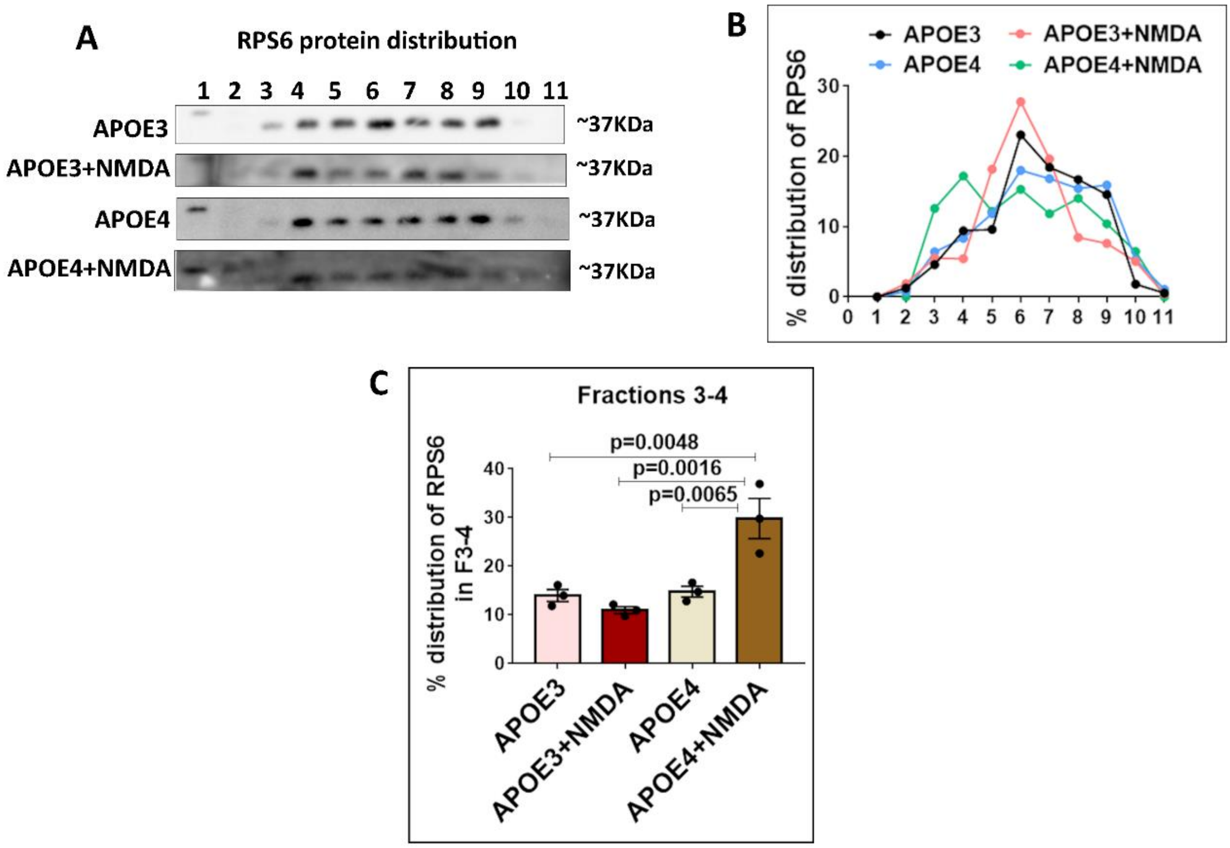


**Supplementary Figure 4 –**

**A -** Rat primary cortical neurons (DIV15) were treated with APOE3/APOE4 (10-15nM) conditioned media for 20 minutes along with NMDAR stimulation for 5 minutes (20µM NMDA). The cell lysates were subjected to polysome profiling and probed for ribosomal protein RPS6. The immunoblots show the distribution of RPS6 on the linear sucrose gradient under different conditions.

**B** –The line graph represents the percentage distribution of RPS6 protein in each fraction under different treatment conditions (quantified from immunoblots of Fig S4A).

**C -** The graph represents the percentage distribution of RPS6 protein in fraction 3-4 under different treatment conditions. Data is represented as mean +/- SEM. N=3, One-way ANOVA ($p=0.0015$) followed by Tukey's multiple comparison test.

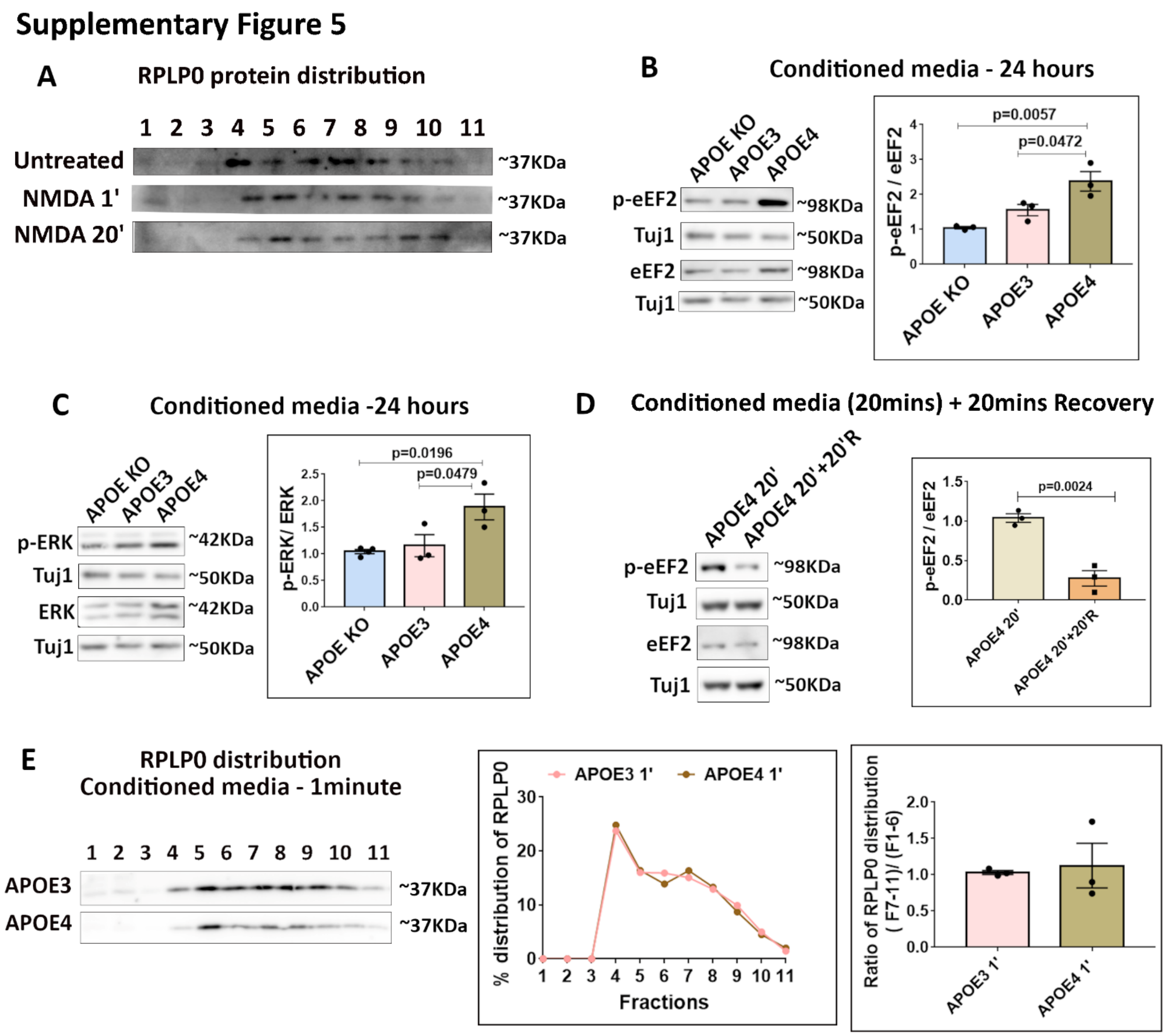
Supplementary Figure 5
A
RPLP0 protein distribution
1 2 3 4 5 6 7 8 9 10 11
Untreated
NMDA 1'
NMDA 20'
~37KDa
B
Conditioned media - 24 hours
APOE KO
APOE3
APOE4
p-eEF2
Tuj1
eEF2
~98KDa
~50KDa
p=0.0057
p=0.0472
p-eEF2 / eEF2
C
Conditioned media -24 hours
p-ERK
ERK
~42KDa
p=0.0196
p=0.0479
p-ERK/ ERK
D
Conditioned media (20mins) + 20mins Recovery
APOE4 20'
APOE4 20'+20'R
p=0.0024
E
RPLP0 distribution
Conditioned media - 1minute
APOE3
APOE4
APOE3 1'
APOE4 1'
% distribution of RPLP0
Fractions
Ratio of RPLP0 distribution
( F7-11)/ (F1-6)

**Supplementary Figure 5 -**

**A** – Rat primary cortical neurons (DIV15) were subjected to NMDAR stimulation for 1 minute and 20 minutes (20µM NMDA). The cell lysates were subjected to polysome profiling and probed for the distribution of ribosomal protein RPLP0. The immunoblots indicate the distribution of RPLP0 in individual fractions.

**B** - Rat primary cortical neurons (DIV15) were treated with APOE (10-15nM) conditioned media for 24 hours and probed for phosphorylation of eEF2. Left - representative immunoblots indicating levels of phospho-eEF2, eEF2 and Tuj1; Right - graph indicating ratio of phospho-eEF2 to eEF2 normalized to Tuj1. Data is represented as mean +/- SEM. N=3, One-way ANOVA (p=0.0068) followed by Tukey's multiple comparison test.

**C** - Rat primary cortical neurons (DIV15) were treated with APOE (10-15nM) conditioned media for 24 hours and probed for phosphorylation of ERK. Left - representative immunoblots indicating levels of phospho-ERK, ERK and Tuj1; Right - graph indicating ratio of phospho-ERK to ERK normalized to Tuj1. Data is represented as mean +/- SEM. N=3, One-way ANOVA (p=0.018) followed by Tukey's multiple comparison test.

**D** - Rat primary cortical neurons (DIV15) were treated with APOE4 (10-15nM) conditioned media for 20 minutes and subjected to recovery for 20 minutes using pre-conditioned neurobasal media. The samples were probed for the phosphorylation of eEF2. Left - representative immunoblots indicating levels of phospho-eEF2, eEF2 and Tuj1; Right - graph indicating ratio of phospho-eEF2 to eEF2 normalized to Tuj1. Data is represented as mean +/- SEM. N=3, Unpaired Student's t-test.

**E** - Rat primary cortical neurons (DIV15) were treated with APOE3/APOE4 conditioned media (10-15nM) for 1 minute. The cell lysates were subjected to polysome profiling and probed for the distribution of ribosomal protein RPLP0. The immunoblots indicate the distribution of RPLP0 in individual fractions. The line graph shows the percentage distribution of RPLP0 under different conditions. The bar graph represents the ratio of RPLP0 protein distribution in fractions 7-11 (translating pool) to fractions 1-6 (non-translating pool).

#### Supplementary Figure 6

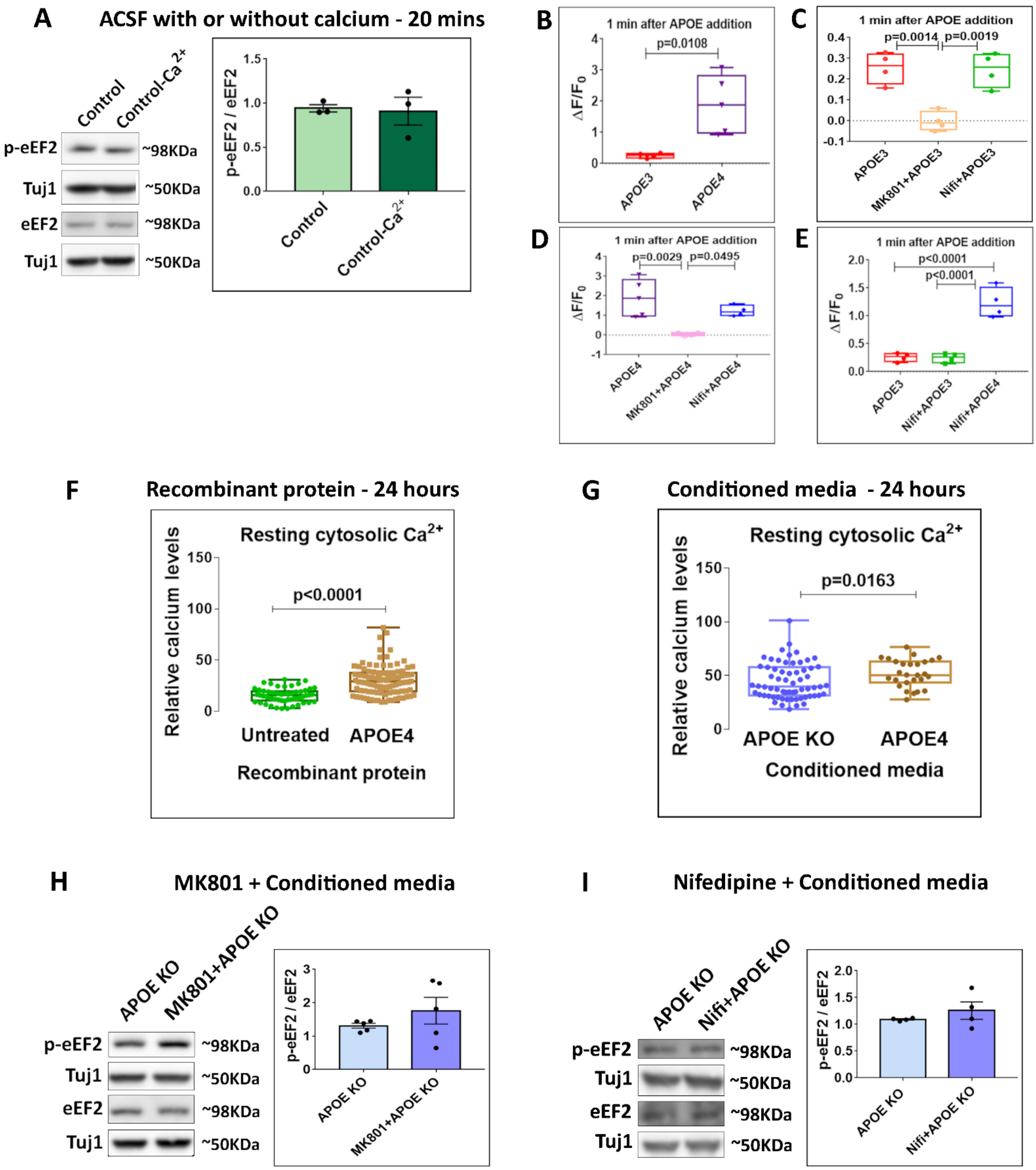

**Supplementary Figure 6 -**

**A** - Rat primary cortical neurons (DIV15) in the presence or absence of extracellular calcium (ACSF with or without calcium) for 20 minutes were probed for phosphorylation of eEF2. Left - representative immunoblots indicating levels of phospho-eEF2, eEF2 and Tuj1; Right - graph indicating ratio of phospho-eEF2 to eEF2 normalized to Tuj1. Data is represented as mean +/- SEM. N=3.

**B, C, D, E** - Box plots represent the quantification of the change in Flou4-AM fluorescence (ΔF/F0) after 1 minute of APOE addition. N=4-5 experiments, each experiment has the average value of 40-50 neurons.

B - Unpaired Student's t-test; C - One-way ANOVA (p=0.0008) followed by Tukey's multiple comparison test; D - One-way ANOVA (p=0.0038) followed by Tukey's multiple comparison test; E - One-way ANOVA (p<0.0001) followed by Tukey's multiple comparison test.

**F** - Rat primary cortical neurons (DIV15) were treated with APOE4 recombinant protein (15nM) for 24 hours and subjected to calcium imaging using Fluo-8AM. The graph shows the resting cytosolic calcium measured in these neurons. N=50-60 neurons from 3 independent experiments. Kolmogorov-Smirnov test.

**G** - Rat primary cortical neurons (DIV15) were treated with APOE KO or APOE4 (10-15nM) conditioned media for 24 hours and subjected to calcium imaging using Fluo-8AM. The graph shows the resting cytosolic calcium measured in these neurons. N=50-60 neurons from 3 independent experiments. Kolmogorov-Smirnov test.

**H** - Rat primary cortical neurons (DIV15) were treated with MK801 (25µM) along with APOE KO conditioned media for 1 minute and probed for phosphorylation of eEF2. Left - representative immunoblots indicating levels of phospho-eEF2, eEF2 and Tuj1; Right - graph indicating ratio of phospho-eEF2 to eEF2 normalized to Tuj1. Data is represented as mean +/- SEM. N=5. One-way ANOVA (ns).

**I** - Rat primary cortical neurons (DIV15) were treated with Nifedipine (50µM) along with APOE KO conditioned media for 20 minutes and probed for phosphorylation of eEF2. Left - representative immunoblots indicating levels of phospho-eEF2, eEF2 and Tuj1; Right - graph indicating ratio of phospho-eEF2 to eEF2 normalized to Tuj1. Data is represented as mean +/- SEM. N=4. One-way ANOVA (ns).
